# Environmental and anthropogenic impacts on key fish and mollusc species of the Mediterranean Sea from the Late Pleistocene until the Industrial Revolution

**DOI:** 10.1101/2025.02.05.636195

**Authors:** Daniela Leal, Konstantina Agiadi, Marta Coll, Maria Bas

## Abstract

Historical baselines are essential for evaluating the cumulative impacts on modern marine ecosystems, particularly in regions such as the Mediterranean, where human activities have been intensive for millennia and climate change is accelerating. However, quantitative evidence of historical impacts remains fragmented. In this study, we investigate changes in the presence, abundance and body size of the Atlantic bluefin tuna, gilthead sea bream, banded-dye murex and turbinate monodont across the Mediterranean Sea from 130,000 years Before Present until the Industrial Revolution (1850 AD), using geological, archaeological, and historical records. Our results reveal significant temporal shifts in the abundance and body size of the banded-dye murex, Atlantic bluefin tuna and gilthead sea bream. Environmental factors, particularly seawater temperature, were the primary drivers of abundance and size in Atlantic bluefin tuna in the past. Human activities, especially exploitation, influenced the abundance of banded-dye murex and Atlantic bluefin tuna, as well as the body size of gilthead sea bream. These findings underscore the importance of integrating long-term ecological data to better understand the interplay between climate, human pressures and ecosystem dynamics.

## 1. INTRODUCTION

Marine ecosystems are major components of the Earth system, supporting biodiversity, human societies and economies (Halpern *et al*. 2008). However, they are increasingly under threat. Understanding how multiple stressors affect marine biota requires long-term ecological records, yet continuous baselines remain scarce (Dietl *et al*. 2015). While most ecological studies rely on short-term datasets (Jackson 1997; Dayton *et al*. 1998), geological, archaeological and historical records offer valuable insights into ecosystems’ long-term ecosystem responses to environmental and anthropogenic pressures (Jackson *et al*. 2001; Lotze *et al*. 2006; Lotze, Coll and Dunne 2011; Fortibuoni *et al*. 2010; Bas *et al*. 2019, 2023; Agiadi and Albano 2020; Agiadi *et al*. 2023, 2024a; Porz *et al*. 2024).

The Mediterranean Sea, a global biodiversity hotspot (IUCN-MED 2009; Coll *et al*. 2010), hosts emblematic species, such as the Atlantic bluefin tuna (*Thunnus thynnus*) (MacKenzie, Mosegaard and Rosenberg 2009) and the Mediterranean monk seal (*Monachus monachus*) (Reijnders, Verriopoulos and Brasseur 1997). Past changes in climate and ocean connectivity have shaped the Mediterranean ecosystems, influencing species migrations and distributions (Agiadi *et al*. 2011, 2018, 2024b). Since the Industrial Revolution, the Mediterranean has warmed faster than the global average (Salat *et al*. 2019; Artana *et al*. Submitted), with profound consequences for marine biodiversity and ecosystems functioning (Garrabou *et al*. 2009; Smale *et al*. 2019; Trisos, Merow and Pigot 2020; Ouled-Cheikh *et al*. 2022).

Marine resources have supported Mediterranean societies for millennia (Madariaga 1964; Mehvar *et al*. 2018). Shellfish exploitation dates back to the Lower Palaeolithic (e.g. ∼300 ka B.P., de Lumley 1966), and by the Bronze Age (5-3 ka BP), extensive fisheries and trade networks were established (e.g., Van Neer *et al*. 2004; Guy *et al*. 2018a; Zohar and Artzy 2019), intensifying throughout the Classical period (Sáez Romero 2014; Mylona 2018). Several key marine species were extensively exploited and/or affected by past environmental changes (Leal, Agiadi and Bas 2025). Long-term exploitation combined with environmental variability, has contributed to the reshaping of marine ecosystems (Lotze *et al*. 2006; Lotze, Coll and Dunne 2011). Today, habitat loss, degradation, and overexploitation continue to alter species productivity, fitness and distribution, with cascading effects on ecosystem structure and functioning (Halpern *et al*. 2008; Coll *et al*. 2010, 2012; Planque *et al*. 2010; Howarth *et al*. 2014; Artana *et al*. Submitted).

In this study, we examine how environmental variability and human activities have influenced commercially relevant species in the Mediterranean Sea over the past 130 kyrs until the Industrial Revolution. Using regional data (Leal, Agiadi and Bas 2025) we analyse Atlantic bluefin tuna (*Thunnus thynnus*), gilthead sea bream (*Sparus aurata*), banded-dye murex (*Hexaplex trunculus*) and turbinate monodont (*Phorcus turbinatus*), testing whether: 1) temperature oscillations from the Late Pleistocene to the Late Holocene (Cacho, Grimalt and Canals 2002; Marchal *et al*. 2002; Rohling *et al*. 2002; Cacho *et al*. 2006; Schmiedl *et al*. 2010) influenced species abundance and body size; and 2) human exploitation compounded these effects over time.

## 2. MATERIALS AND METHODS

### 2.1 Study area

The Mediterranean Sea is a semi-enclosed basin (3,000,000 km^2^, maximum depth ∼5,300 m (Robinson and Golnaraghi 1994). For analytical purposes, we divided the basin into three sub-regions: Western, Eastern and Adriatic, defined by the straits of Sicily and Otranto (Millot and Taupier-Letage 2005; Rio *et al*. 2007; Robinson *et al*. 2009) (Figure 1). The Marmara and Black seas were included in the Eastern Mediterranean.

**Figure 1-.**
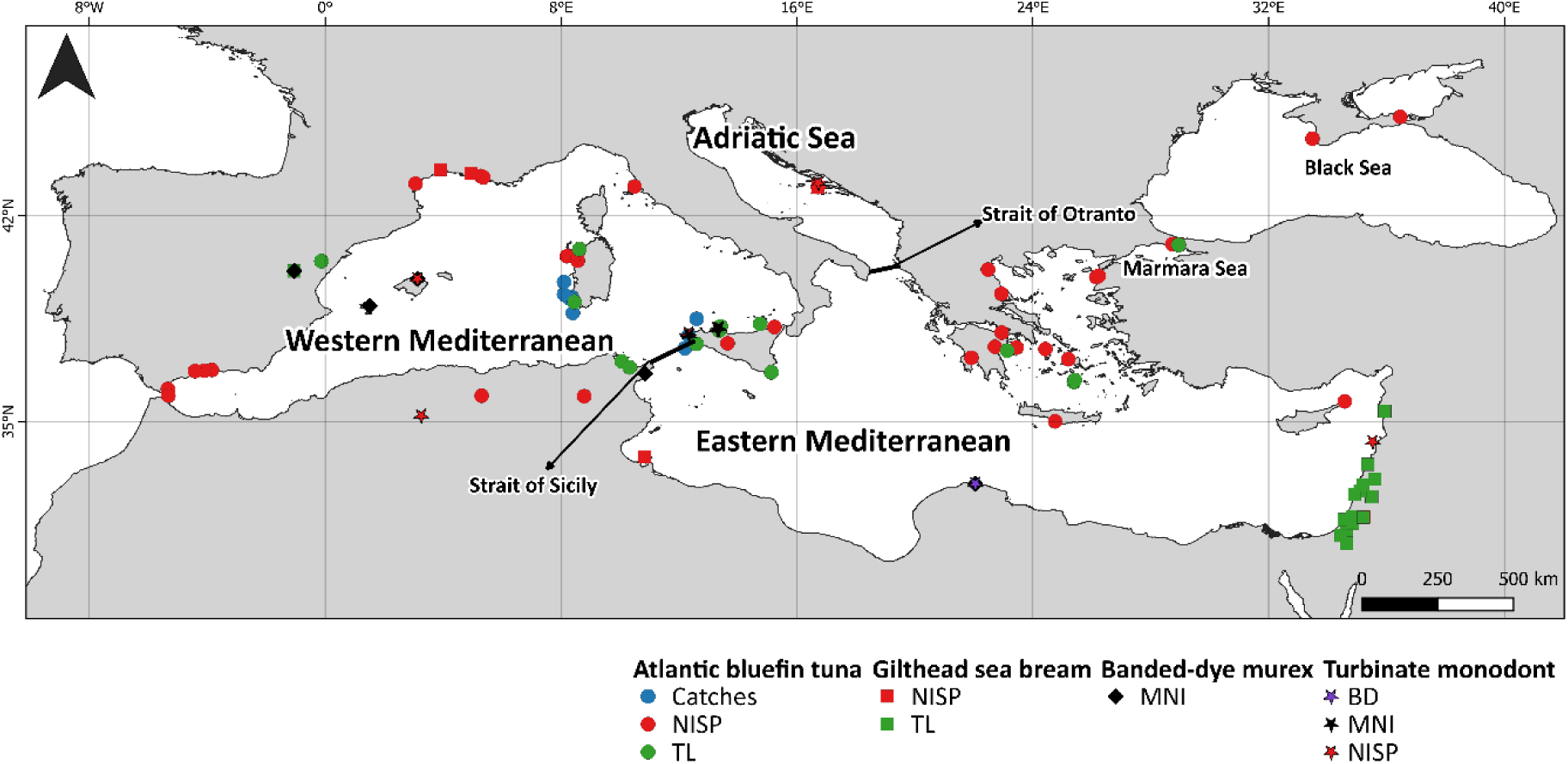
Map of the Mediterranean Sea showing the sub-regions (Western Mediterranean, Eastern Mediterranean and Adriatic Sea) and their natural divisions (Strait of Otranto and Strait of Sicily), with the data locations (Table S6). Variables used include the Number of Identified Specimens (NISP). Minimum Number of Individuals (MNI), Total Length (TL) and Basal Diameter (BD).

### 2.2 Species and temporal framework

Species data were available across both spatial and temporal scales within the Mediterranean sub-regions, although catch data were only available for the Atlantic bluefin tuna.

We adopted the temporal divisions defined by the International Commission on Stratigraphy (ICS; Cohen, Harper and Gibbard 2013) into the Late Pleistocene (130,000–11,650 years Before Present (BP); Head *et al*. 2021), Early Holocene (11,650–8,186 years BP; Stuiver and Reimer 1986), Middle Holocene (8,186–4,200 years BP; Stuiver and Reimer 1986) and Late Holocene (4,200–0 years BP; Stuiver and Reimer 1986). Each period is characterised by major climatic events (e.g., glacial-interglacial cycles, Lionello *et al*. 2023) and societal transitions (e.g., from hunter-gatherers to sedentary communities, Isern *et al*. 2014; Valla 2018). The Late Pleistocene includes the Last Interglacial (130-116 kyrs BP; Kukla *et al*. 2002) and the Younger Dryas (14,749–13,646 years BP; Benjamin *et al*. 2017). The Holocene features notable temperature fluctuations, such as the Medieval Warm Period (MWP; 1,050–650 BP) and the Little Ice Age (LIA; 650–100 years BP) (Cisneros *et al*. 2016).

Mediterranean fisheries evolved significantly between the pre-industrial (130 kyrs - 1850 AD) to the post-industrial era (1850 AD – present day). Fishing practices transitioned from manual and artisanal techniques (harpoons, gillnets and traps; Ruas 2005; Roberts *et al*. 2011; Horden and Purcell 2013; Pedergnana *et al*. 2021), to mechanised fishing and steam-powered trawlers with the onset of industrial fishing (1880 AD) (Gabriel *et al*. 2005).

### 2.3 Database

The database compiled by Leal, Agiadi and Bas (2025a) was developed through a systematic review following the PRISMA methodology (Page *et al*. 2021), integrating peer-reviewed and secondary literature with geological, archaeological and historical sources (Table S7). Presence and abundance data were quantified using two zooarchaeological metrics, which are commonly used to infer patterns of human exploitation of faunal resources: the Number of Identified Specimens (NISP; Grayson 1984) and Minimum Number of Individuals (MNI; Lyman 1994).. The review also incorporated over 1,200 entries of Atlantic bluefin tuna catch data from Italian tuna traps (1700–1936; Addis *et al*. 2008; Longo and Clark 2012; Polanco-Martínez *et al*. 2018), which serve as long-term proxies for abundance (Ravier 2001). Historical abundance trends were analysed in relation to Sea Surface Temperature (SST; Jalali *et al*. 2016), human civilisations and fishing technologies. Body size (BS) data were derived from multiple sources: a) Total Length (TL) for Atlantic bluefin tuna from Andrews *et al*. (2022, 2023a, 2023c, 2023b); b) TL for gilthead sea bream from Western archaeological sites (Fernández-López De Pablo and Gabriel 2016) and Eastern sites (Guy *et al*. 2018; Fuller *et al*. 2020); and c) Basal Diameter (BD) of the turbinate monodont from Haua Fteah cave, Libya (Hunt *et al*. 2011; Prendergast *et al*. 2016).

### 2.4 Data Analyses

Normality and homoscedasticity were tested using the Shapiro–Wilk and Levene’s tests, respectively (*car* R package; Fox and Weisberg 2019). For NISP and MNI, non-parametric tests were applied: the Mann-Whitney U test for two-group comparisons and the Kruskal-Wallis test for multi-group comparisons, followed by post-hoc pairwise Wilcoxon tests. For TL, a two-sample t-test for Western Mediterranean tuna and a Mann-Whitney *U* test for the Eastern Mediterranean tuna. For the gilthead sea bream, ANOVA with a Tukey’s HSD post-hoc test was used to compare three periods. For BD, ANOVA (mean values) and Welch ANOVA (maximum values) were applied to multi-period comparisons. Catch data trends were analysed using the Mann-Kendall test (*trend* R package; Pohlert 2015). Data classification was performed using Jenks’ natural breaks method (Jenks 1977) (*classInt* R package; Bivand 2006). Inter-period differences were assessed with Kruskal-Wallis and pairwise Wilcoxon tests. A significance threshold of *p* ≤ 0.05 was applied, with Bonferroni correction. Maps were produced in QGIS 3.34.13, and statistical analyses in R 4.4.3 (R Core Team 2025).

## 3. RESULTS

### 3.1 Presence and abundance of targeted species

NISP data for gilthead sea bream and Atlantic bluefin tuna span the entire temporal framework. However, comparisons between periods within each region revealed no statistically significant differences (Table S2.1; Figures S3.1B–S3.4B). Among molluscs, only turbinate monodont presented sufficient data for temporal analysis across the Mediterranean Sea, but no significant differences were detected across periods (Table S2.1; Figures S3.5B and S3.6B).

MNI data for turbinate monodont were available for both the Western and Eastern Mediterranean, yet no significant differences were found between periods (Table S2.2, Figures S3.7 and S3.8). In contrast, banded-dye murex showed significant temporal variation in the MNI in the Western Mediterranean (*H_2_* = 15.792; *p* = 0.0004, Figure 2A), particularly between the Early and Late Holocene (*p* = 0.0013; Figure 2B; Table S2.2).

**Figure 2-.**
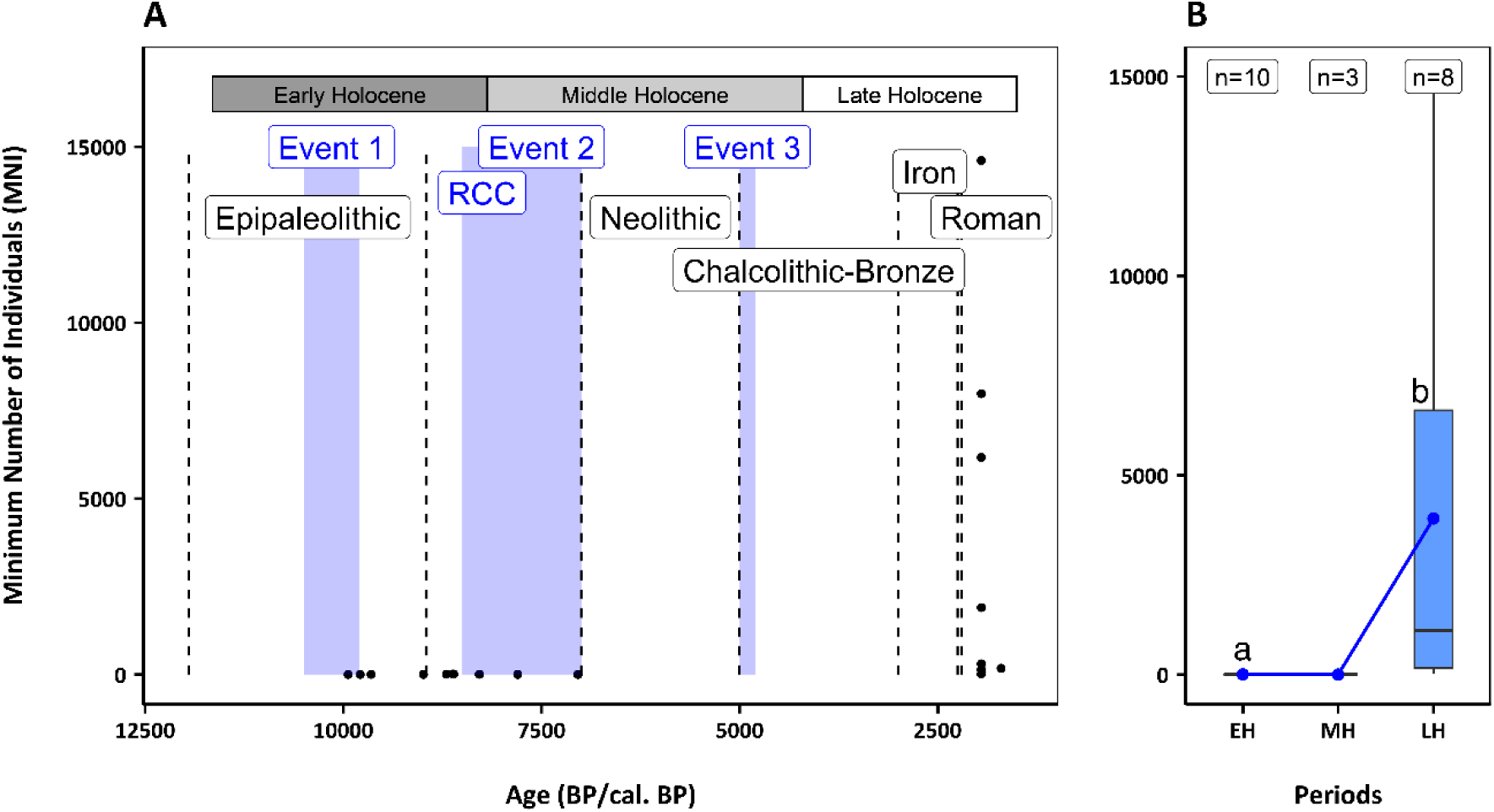
Distribution of Minimum Number of Individuals (MNI) for banded-dye murex in the Western Mediterranean. (A) Temporal distribution of records, with major climatic events (purple rectangles) and human cultural periods (black dashed lines); (B) Boxplots of MNI data by geological period: Early Holocene (EH), Middle Holocene (MH), and Late Holocene (LH). Blue dots represent mean values and bars indicate standard deviation. Letters denote statistically significant differences. Sample sizes (n) are shown above each boxplot.

### 3.2 Catch records

Catch data (annual totals) were available only for Atlantic bluefin tuna from 250 to 14 years BP, corresponding to the period between 1700 and 1936 AD (Figure 3). Recorded values ranged from 0 to approximately 12,500 individuals per year along the coasts of Sardinia and Sicily in the Western Mediterranean (Figure 1). Two major peaks were observed: around 200 years BP (∼1750 AD) during the LIA with ∼10,000 individuals, and between 75–45 years BP (1875–1905 AD), following the Industrial Revolution (1850 AD) (∼12,000 individuals).

**Figure 3-.**
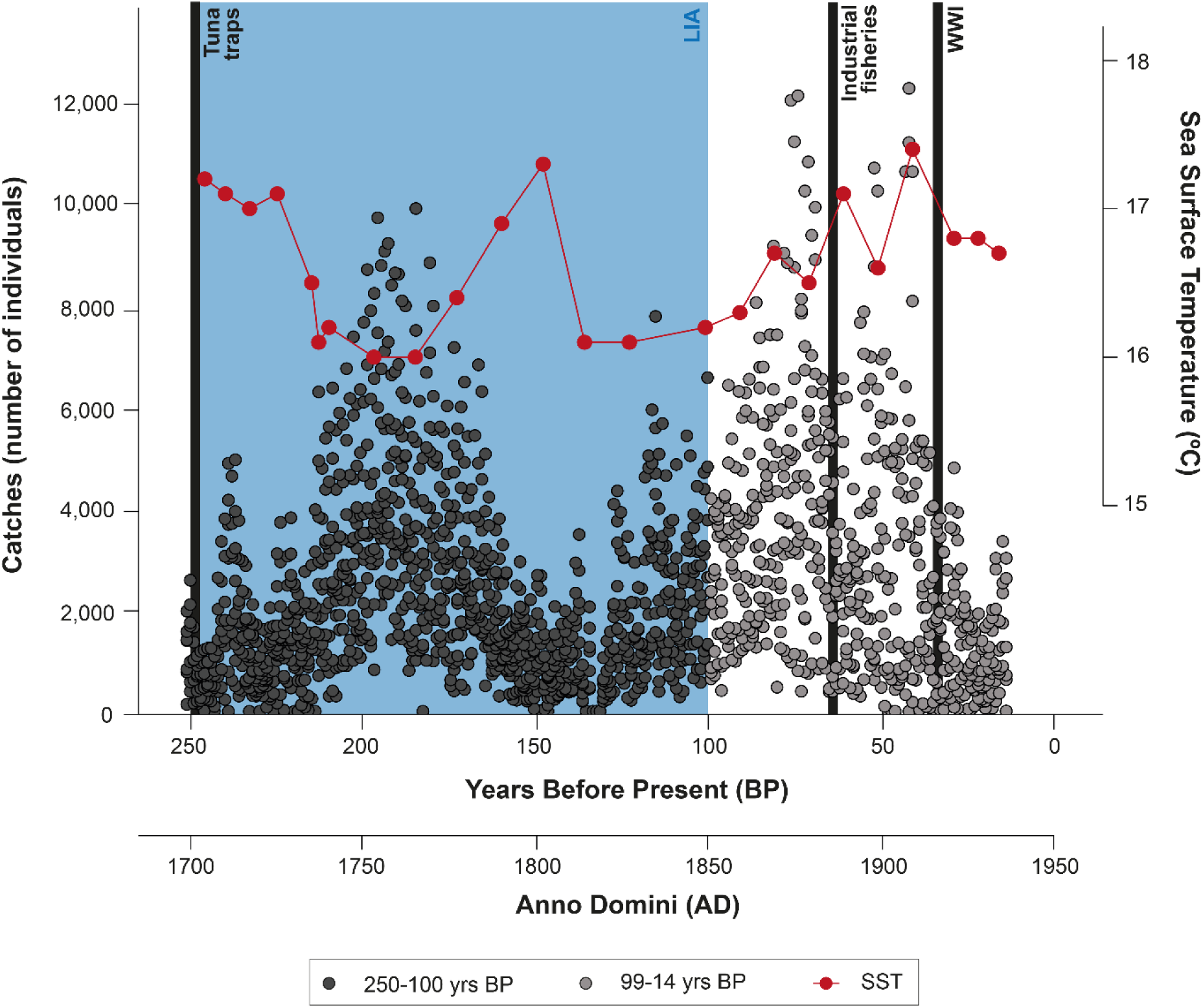
Atlantic bluefin tuna catch records from the Late Holocene (250–14 years BP;1700–1936 AD), based on tuna traps from the Western Mediterranean Sea. Black points represent pre-industrial catches (250–100 years BP; 1700–1850 AD), grey points represent post-industrial catches (99–14 years BP; 1851–1936 AD). Red points indicate reconstructed Sea Surface Temperature (SST) data from the Gulf of Lions (Jalali *et al*. 2016). The blue rectangle refers to the Little Ice Age (LIA). Vertical lines indicate the use of tuna traps, the onset of industrial fisheries (70 BP; 1880 AD; von Brandt 1972; Gabriel *et al*. 2005) and WWI (World War I). Note: tuna trap technology remained largely unchanged until the 1960s (Doumenge 1998).

A significant declining trend in catches was detected between 250-14 years BP (*S* = −3.98 e^5^; Kendall’s tau coefficient = −2.39 e^−1^; *p* < 2.2e^−16^), with differences between specific intervals (*H_5_* = 407.55; *p* < 2.2e^−16^; Table S4; Figure S4).

### 3.3 Body size

In the Western Mediterranean, a significant increase in the TL of Atlantic bluefin was found between the MWP and the LIA (*t_15.141_* = 2.967; *p* = 0.01; Figures 4 and S5.1; Table S2.4).

**Figure 4-.**
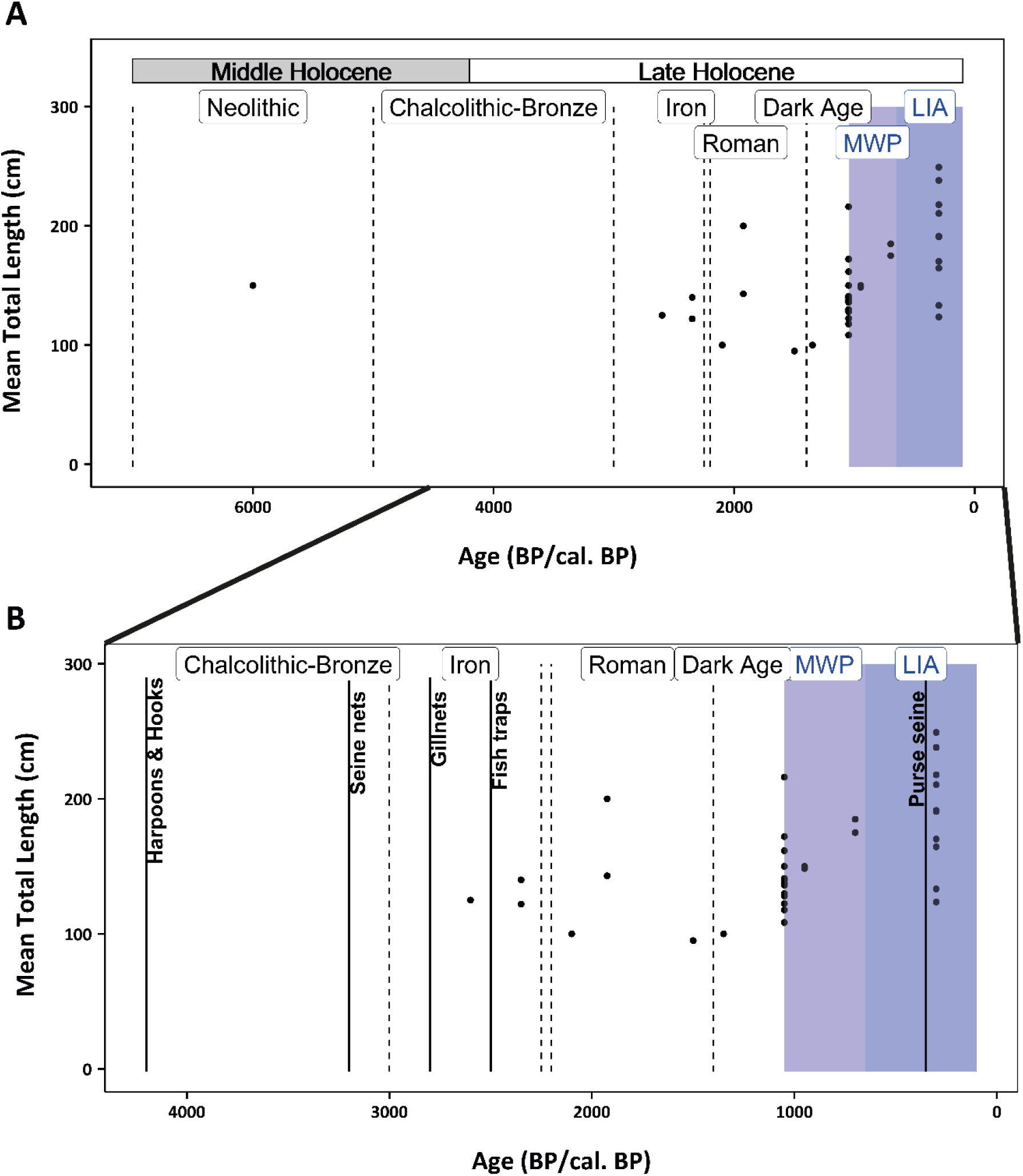
Distribution of Total Length (TL) data for Atlantic bluefin tuna in the Western Mediterranean. (A) Temporal distribution of records, including major climatic events (purple rectangles) and anthropogenic periods (black dashed lines); (B) Late Holocene distribution of records, with vertical lines indicating the main fishing technologies used during the period.

In the Eastern Mediterranean, TL of Atlantic bluefin tuna also showed significant increases between the Roman Period (2,500–1,050 years BP) and the MWP (1,050–650 years BP), as well as within the MWP itself (*W* = 8; *p* = 0.01035) (Figures 5 and S5.2; Table S2.4).

**Figure 5-.**
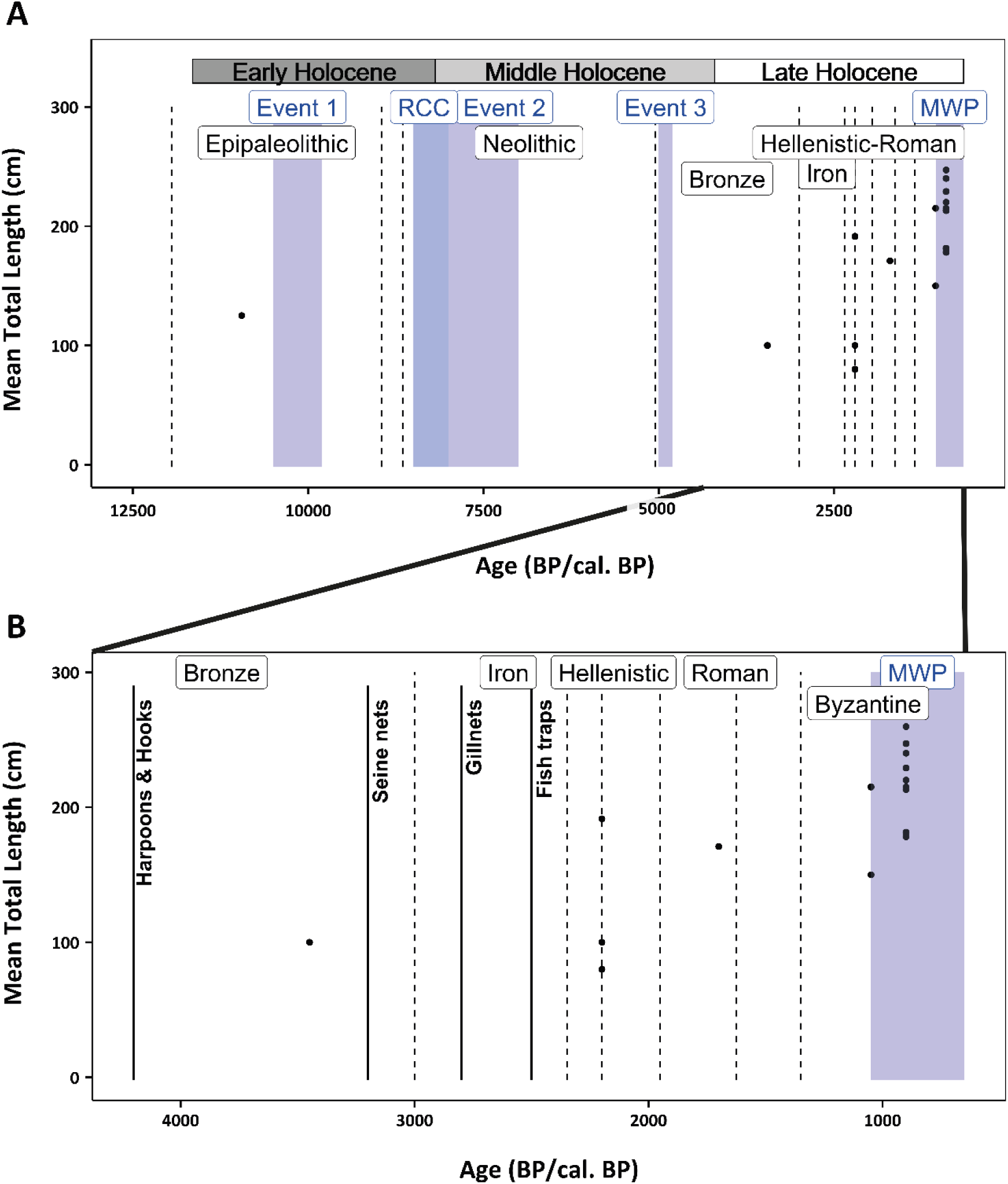
Distribution of Total Length (TL) data for Atlantic bluefin tuna in the Eastern Mediterranean. (A) Temporal distribution of records, including major climatic events (purple rectangles) and anthropogenic periods (black dashed lines); (B) Late Holocene distribution of records, with vertical lines indicating the main fishing technologies used during the period.

For gilthead sea bream, TL data were available for the Eastern Mediterranean (Figure 6). Pairwise comparisons (F = 10.28; *p* = 7.54e-^5^) revealed significant reductions in TL between the Early and Late Holocene (*p* = 0.0061), the Middle and Late Holocene (*p* = 0.0010; Figure 6B), and between the Bronze and Iron Ages and the Byzantine Period (*H_2_* = 8.501; *p* = 0.014, Table S2.4 and Figure S5.3).

**Figure 6-.**
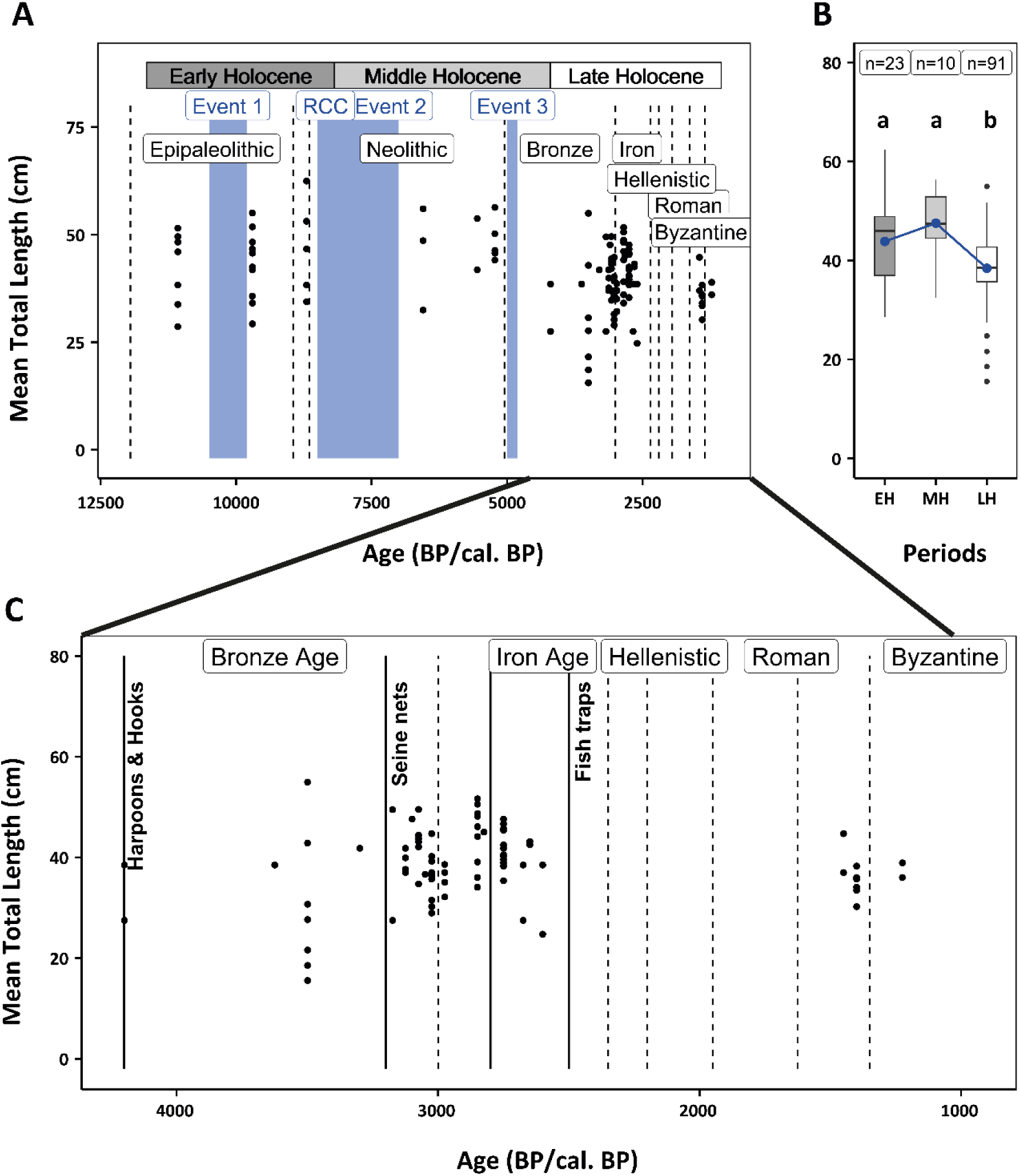
Distribution of Total Length (TL) data for gilthead sea bream in the Eastern Mediterranean. (A) Temporal distribution of records, including major climatic events (purple rectangles) and anthropogenic periods (black dashed lines); (B) Boxplots of TL data for each geological period: Early Holocene (EH), Middle Holocene (MH), and Late Holocene (LH). Blue dots represent mean values, and bars indicate standard deviation. Letters denote statistically significant differences. Sample sizes (n) are shown above each boxplot; (C) Late Holocene distribution of records, with vertical lines indicating the main fishing technologies used during the period.

For the turbinate monodont, BD data were available for the Eastern Mediterranean, including both mean and maximum values. No significant differences were found between the geological periods, for either metric (Table S2.5; Figures S5.4-S5.5).

## 4. DISCUSSION

### 4.1 Environmental Driven Changes in Atlantic Bluefin Tuna

Atlantic bluefin tuna is a highly migratory species that travels between cold feeding grounds in the North Atlantic and warm spawning areas in the Mediterranean Sea (Mather, Mason and Jones 1995; Block *et al*. 2001, 2005; Rooker *et al*. 2007). Climatic changes influence its growth, reproduction, population size, mortality and migration patterns (Pepin 1991; Graham and Dickson 2004). Our data show a peak in tuna catches during the late LIA, which may be linked to cooler conditions (Cisneros *et al*. 2016; Margaritelli *et al*. 2020) and enhanced nutrient availability and primary productivity in the North Atlantic (Pourmand *et al*. 2007), resulting in a greater number of mature individuals entering the Mediterranean. Additionally, tuna migration may have shifted toward the basin during the LIA, due to increased climatic instability and extreme events, leading to higher recruitment in the Western Mediterranean and greater concentration of schools near historical tuna traps (Ravier and Fromentin 2004, Ganzedo *et al*. 2009). Considering the 6–7 year lag between hatching and recruitment (Zorita *et al*. 2005), the increased landings observed in the later stages of the LIA may reflect changes in distribution and behaviour rather than an overall increase in population size.

Temperature also plays a key role in shaping plankton dynamics, influencing both production and distribution (e.g., McGowan, Cayan and Dorman 1998; Beaugrand *et al*. 2002). Our results suggest that cooler conditions during the late LIA enhanced trophic productivity, increasing tuna availability and catch rates in the Western Mediterranean. Similar patterns have been observed for Pacific bluefin tuna, where SST is a stronger driver of recruitment than spawning stock biomass, likely due to its influence on juvenile habitat suitability (Muhling *et al*. 2018). Temperature-driven changes in plankton propagate through the food web, altering prey availability and influencing tuna migration (Mather, Mason and Jones 1995; Ravier and Fromentin 2004).

We also observed an increase in TL of Atlantic bluefin tuna during the late LIA, likely driven by cooler conditions, improved feeding opportunities, and altered migration patterns that concentrated larger individuals in the region. This aligns with the species’ recruitment lag (Santamaria *et al*. 2009), explaining the delayed increase in adult abundance and TL. Although warmer conditions can enhance consumer-resource interactions (Allen, Gillooly and Brown 2005; López-Urrutia *et al*. 2006; O’Connor 2009), the increase in TL under cooler SST suggests indirect effects, such as enhanced trophic productivity or greater adult retention, played a key role. The species’ broad dietary niche (Olesen and Jordano 2002; Pardo *et al*. 2025) likely supported adaptation to shifting prey fields, while cooler, more productive conditions may have improved larval growth and survival (García *et al*. 2006; Catalán *et al*. 2011; Reglero *et al*. 2018; Trueman *et al*. 2023). Consistent with other Mediterranean fish (Hattab *et al*. 2021), larger body sizes under colder temperatures indicate that long-lived top predators like the Atlantic bluefin tuna (Santamaria *et al*. 2009) may deviate from general patterns due to their complex ecological responses (Fromentin and Powers 2005).

### 4.2 Human Exploitation and Species Decline

This study highlights the long-term effects of human exploitation on species abundance and body size in the Mediterranean Sea, supported by historical and ethnographic records of traditional practices (Galili, Zemer and Rosen 2013; De Nicolò 2018; Lucchetti *et al*. 2023).

The banded-dye murex was intensively harvested in the Late Holocene, particularly in the Roman Period (Oliver 2015). Its intertidal habitat made it easily accessible to early communities (Bonanno *et al*. 2016; Fernández-López De Pablo and Gabriel 2016) and later enabled large-scale exploitation for purple dye production (Michel and McGovern 1987; Oliver 2015). Shellfish provided a stable, year-round food resource, supporting coastal populations during periods of environmental stress or resource scarcity (Erlandson 1988; Colonese *et al*. 2011). However, this accessibility also made them vulnerable to overexploitation, especially with the rise of industrial dye production (Michel and McGovern 1987; Erlandson *et al*. 2008; Oliver 2015). These dynamics illustrate the dual role of shellfish as both resilient resources and indicators of human-induced ecosystem change.

The long-term decline in Atlantic bluefin tuna abundance reflects sustained fishing pressure (Polanco-Martínez *et al*. 2018; this study). Traditional tuna traps, once considered sustainable (De la Serna *et al*. 2012; Longo 2012), were eventually replaced by industrial technologies (e.g., iron anchors, steel cables; Lentini 2001), and steam-powered gears; Joseph 2003), which enabled large-scale exploitation and contributed to the stock collapse (Andrews *et al*. 2022). Removing top predators can destabilise food webs through top-down effects (Pauly *et al*. 1998; Scheffer, Carpenter and Young 2005). Although direct evidence of trophic cascades involving tuna in the Mediterranean Sea is limited, isotopic and archaeological data suggest shifts in its trophic ecology during historical periods (Andrews *et al*. 2022, 2023c), altering predator-prey dynamics. Ecopath models from the Catalan Sea confirm these findings, showing food web restructuring following the loss of top and intermediate trophic-level species (Forrestal *et al*. 2012), consistent with broader patterns in the Mediterranean (Stergiou, Tsikliras and Pauly 2009) and the North Sea (Mariani *et al*. 2017).

### 4.3 Changes in Body Size

Overexploitation may have contributed to the reduction in body size of gilthead sea bream from the Early to the Late Holocene, particularly between the Bronze and Iron Age and the Byzantine Period. In contrast, Atlantic bluefin tuna showed the opposite trend, with increasing body sizes from the Roman Period through the MWP and the LIA.

Historical records show declining tuna landings in the Gulf of Cadiz between the mid-16^th^ and 18^th^ centuries (Sarmiento 1757), likely due to juvenile overfishing and habitat degradation (García 2016). However, tuna body size increased during the MWP and LIA, possibly reflecting fisheries-induced evolution (FIE), where intense exploitation selects for early-maturing, faster-growing genotypes (Law 2000; Enberg *et al*. 2012). Similar long-term effects have been documented in Baltic cod, with larger individuals in the Neolithic (∼4500 yrs BP) compared to modern specimens (Limburg *et al*. 2008). From the Iron Age onward, the adoption of fish traps (Mylona 2021) and later industrial gears such as purse seines marked a shift to large-scale fisheries, likely influencing tuna body size across the basin.

Gilthead sea bream showed signs of overexploitation in the Eastern Mediterranean during the Late Holocene, especially from the Bronze Age to the Byzantine period, with a marked decline in body size (Guy *et al*. 2018; this study). Similar trends have been observed in other sparids, such as *Archosargus probatocephalus,* under long-term fishing pressure (Guiry *et al*. 2021). Zooarchaeological records from Orkney (Scotland) also reveal offshore overfishing of large cod between the 11^th^ and 13^th^ centuries, followed by declines in abundance and size (Harland and Barrett 2012). The shift to industrial-scale, less selective fishing gear (Galili, Zemer and Rosen 2013) likely reinforced these reductions. Size-selective harvesting can alter trophic interactions and reduce ecosystem resilience (Shackell *et al*. 2010; Garcia *et al*. 2012), underscoring the long-term ecological consequences of sustained exploitation.

### 4.4 Limitations

While this study provides valuable insights, several limitations must be acknowledged. Our analyses focused on data-rich, historically and economically relevant species, potentially overlooking less-studied taxa and broader ecosystem dynamics. Data scarcity, particularly in the Adriatic Sea, reflects the uneven availability of published records (Leal, Agiadi and Bas 2025). Most data originate from coastal archaeological sites, which may bias the representation of past exploitation patterns (Galili and Rosen 2008; Leal, Agiadi and Bas 2025). Additional constraints include variability in dating, body size measurements, and taphonomic preservation, which limit the accuracy of past population reconstructions and inter-period comparability (Agiadi *et al*. 2024b, 2024a). Moreover, uncertainty regarding the fishing gear used and the exclusion of non-peer-reviewed sources, such as logbooks or diaries, introduce further biases (Agiadi *et al*. 2024a). Despite these limitations, our study advances the understanding of how environmental variability and human activities have shaped key Mediterranean marine species over long timescales. Emerging techniques, such as ancient (aDNA) and sedimentary ancient DNA (*seda*DNA), offer promising avenues for higher resolution reconstructions of paleoecological conditions and ecosystem dynamics (De Schepper *et al*. 2019; Monk *et al*. 2021; Kjær *et al*. 2022).

## Conclusions

This study underscores the importance of adopting long-term perspectives to understand changes in the marine ecosystem. Our findings suggest that both environmental and human factors influenced Atlantic bluefin tuna, while banded-dye murex and gilthead sea bream were mainly affected by sustained human pressure. Understanding these historical trajectories helps clarify the mechanisms and magnitude of ecological shifts, contributing to the interpretation of “shifting baselines” (Pauly *et al*. 1998). By integrating palaeoecological, archaeological, and historical records, this research provides a robust framework for reconstructing historical baselines and informing future management of Mediterranean marine ecosystems.

## ACKNOWLEDGEMENTS

This research contributes to the objectives of Q-MARE (a PAGES working group), and it is part of the Integrated Marine Ecosystem Assessments (iMARES) research group funded by Agència de Gestió d’Ajuts Universitaris i de Recerca (Generalitat de Catalunya).

## AUTHOR CONTRIBUTIONS

Conceptualisation: DL, KA, and MB. Methodology: DL, KA, MC and MB. Data curation: DL. Formal analysis: DL. Writing—original draft: DL. Writing—review & editing: KA, MC and MB. Supervision: KA and MB.

## DECLARATION OF INTERESTS

The authors declare that there are no conflicts of interest.

## FUNDING

This work was supported by the Erasmus+ Program Grant no. 2022-1-PT01-KA131-HED-000061155; MCIN/AEI/10.13039/501100011033 and NextGenerationEU/PRTR, Grant no. FJC2020-043762-I; EU HORIZON-CL6-2021-BIODIV-01-04 project Ges4Seas (Grant Agreement 101059877); the “Severo Ochoa Centre of Excellence” accreditation grant CEX2024-001494-S funded by AEI 10.13039/501100011033.

## DATA AVAILABILITY

All data produced for this work are available within the manuscript and in the supplementary material.

## Supplementary material

### S1. List of environmental and cultural events

**Table S1-.**
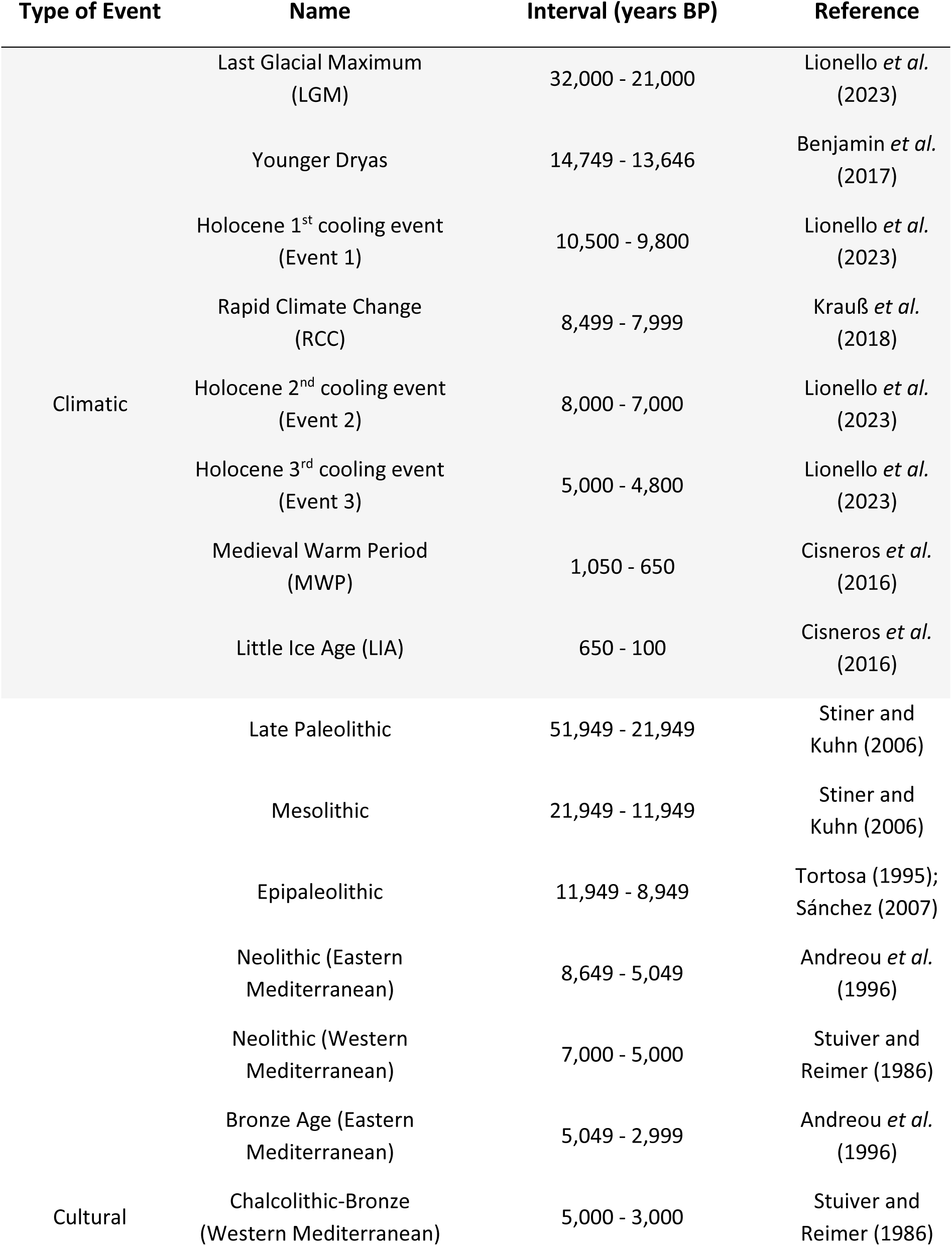
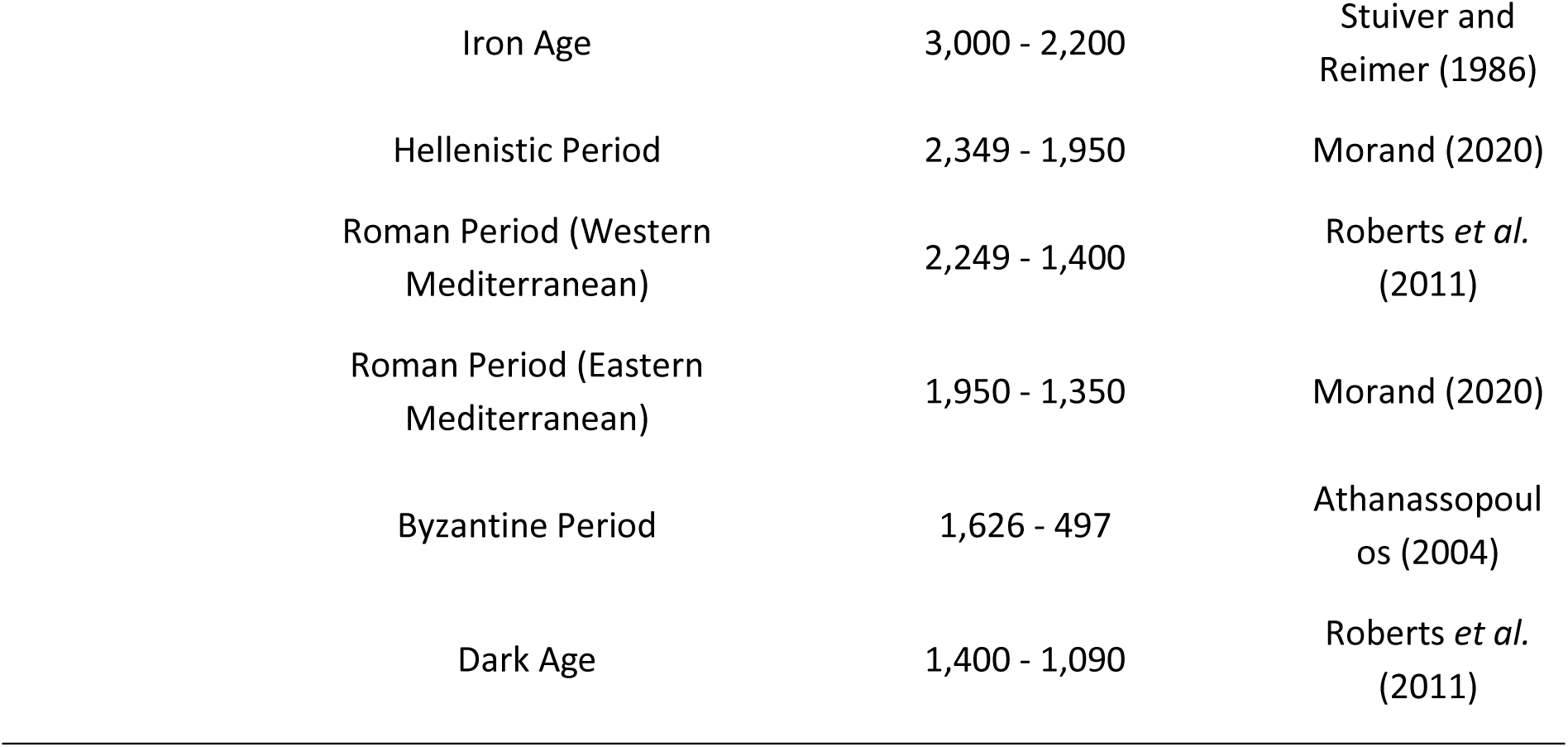
List of environmental and cultural events, with corresponding names, time intervals and references.

### S2. Results from statistical analyses

**Table S2.1-.** Results from the statistical analyses performed regarding NISP data for the Atlantic bluefin tuna, the gilthead sea bream and the turbinate monodont by sub-regions of the Mediterranean Sea.

| Tests | Shapiro-Wilk test |  | Levene's test |  | Kruskal-Wallis |  | Mann-Whitney *U* |
| --- | --- | --- | --- | --- | --- | --- | --- |
| Species/<br>Sub-region | Western | Eastern | Western | Eastern | Western | Eastern | Western |
| Atlantic bluefin tuna | $p = 2.2e^{-16}$ | $p = 4.917e^{-15}$ | $p = 0.917$ | $p = 4.99e^{-15}$ | $H_2 = 0.0504$<br>$p = 0.9751$ | $H_2 = 8.1208$<br>$p = 0.01724$ | |
| Gilthead sea bream | $p = 2.2e^{-16}$ | $p = 2.2e^{-16}$ | $p = 0.5199$ | $p = 0.9085$ | | $H_2 = 1.636$<br>$p = 0.4413$ | $W = 108.5$<br>$p = 0.866$ |
| Turbinate monodont | $p = 9.22e^{-6}$ | $p = 6.454e^{-5}$ | $p = 0.6171$ | $p = 0.6171$ | $H_3 = 4.5617$<br>$p = 0.2069$ | $H_2 = 0.0362$<br>$p = 0.982$ | |

**Table S2.2-.** Results from the statistical analyses performed regarding MNI data for the banded-dye murex for the Western Mediterranean. Numbers in bold represent statistical differences.

| Tests | Shapiro-Wilk test |  | Levene's test |  | Kruskal-Wallis |  |
| --- | --- | --- | --- | --- | --- | --- |
| Species/<br>Sub-region | Western | Eastern | Western | Eastern | Western | Eastern |
| Banded-dye murex | $W = 0.480$<br>$p = 1.349e^{-7}$ | | $F = 4.270$<br>$p = 0.03$ | | $H_2 = 15.792$<br>**$p = 0.0004$** | |
| Turbinate monodont | $p = 2.247e^{-7}$ | $p = 9.57e^{-5}$ | $p = 0.5961$ | $p = 0.3724$ | $H_3 = 0.8223$<br>$p = 0.8441$ | $H_2 = 1.667$<br>$p = 0.4346$ |

**Table S2.3-.** Results from the statistical analyses performed regarding catch data for the Atlantic bluefin tuna for the Western Mediterranean, for periods divided using the Jenks natural breaks classification method (*classInt* package; Bivand 2023).

| Tests | Shapiro-Wilk test | Levene's test | Kruskal-Wallis test |
| --- | --- | --- | --- |
| Atlantic bluefin tuna | $W = 0.91514$<br>$p = 3.12e^{-12}$ | $F = 33.624$<br>$p < 2.2e^{-16}$ | $H_5 = 407.55$<br>**$p < 2.2e^{-16}$** |

**Table S2.4-.** Results from the statistical analyses performed regarding total length (TL) data for the Atlantic bluefin tuna and the gilthead sea bream by sub-regions of the Mediterranean Sea.

| Tests<br>Species/<br>Sub-region | Shapiro-Wilk test |  | Levene's test |  | Two-sample t-test | Mann-Whitney<br>U test | one-way<br>ANOVA |
| --- | --- | --- | --- | --- | --- | --- | --- |
|  | Western | Eastern | Western | Eastern | Western | Eastern | Eastern |
| Atlantic bluefin tuna | $W = 0.9593$<br>$p = 0.1686$ | $W = 0.9091$<br>$p = 0.0163$ | $F = 2.103$<br>$p = 0.1554$ | $F = 0.9167$<br>$p = 0.3468$ | $t = 2.967$<br>$p = \mathbf{0.01}$ | $W = 8$<br>$p = \mathbf{0.01035}$ | |
| Gilthead sea bream | | $W = 0.9898$<br>$p = 0.4921$ | | $F = 1.5013$<br>$p = 0.227$ | | | $F = 10.28$<br>$p = \mathbf{7.54e^{-05}}$ |

**Table S2.5-.** Results from the statistical analyses performed regarding basal diameter (BD) data (mean and maximum values) for the Turbinate monodont for the Eastern Mediterranean.

| Tests<br>Variable/<br>Sub-region | Shapiro-Wilk test |  | Levene's test |  | one-way<br>ANOVA | Welch<br>ANOVA |
| --- | --- | --- | --- | --- | --- | --- |
|  | Mean | Maximum | Mean | Maximum | Mean | Maximum |
| Turbinate monodont | $W = 0.9102$<br>$p = 0.245$ | $W = 0.9794$<br>$p = 0.4367$ | $F = 1.2519$<br>$p = 0.3364$ | $F = 4.4836$<br>$p = 0.0071$ | $F = 2.021$<br>$p = 0.195$ | $F = 0.3701$<br>$p = 0.7767$ |

### S3. Presence and abundance of the targeted species

**Figure S3.1-.**
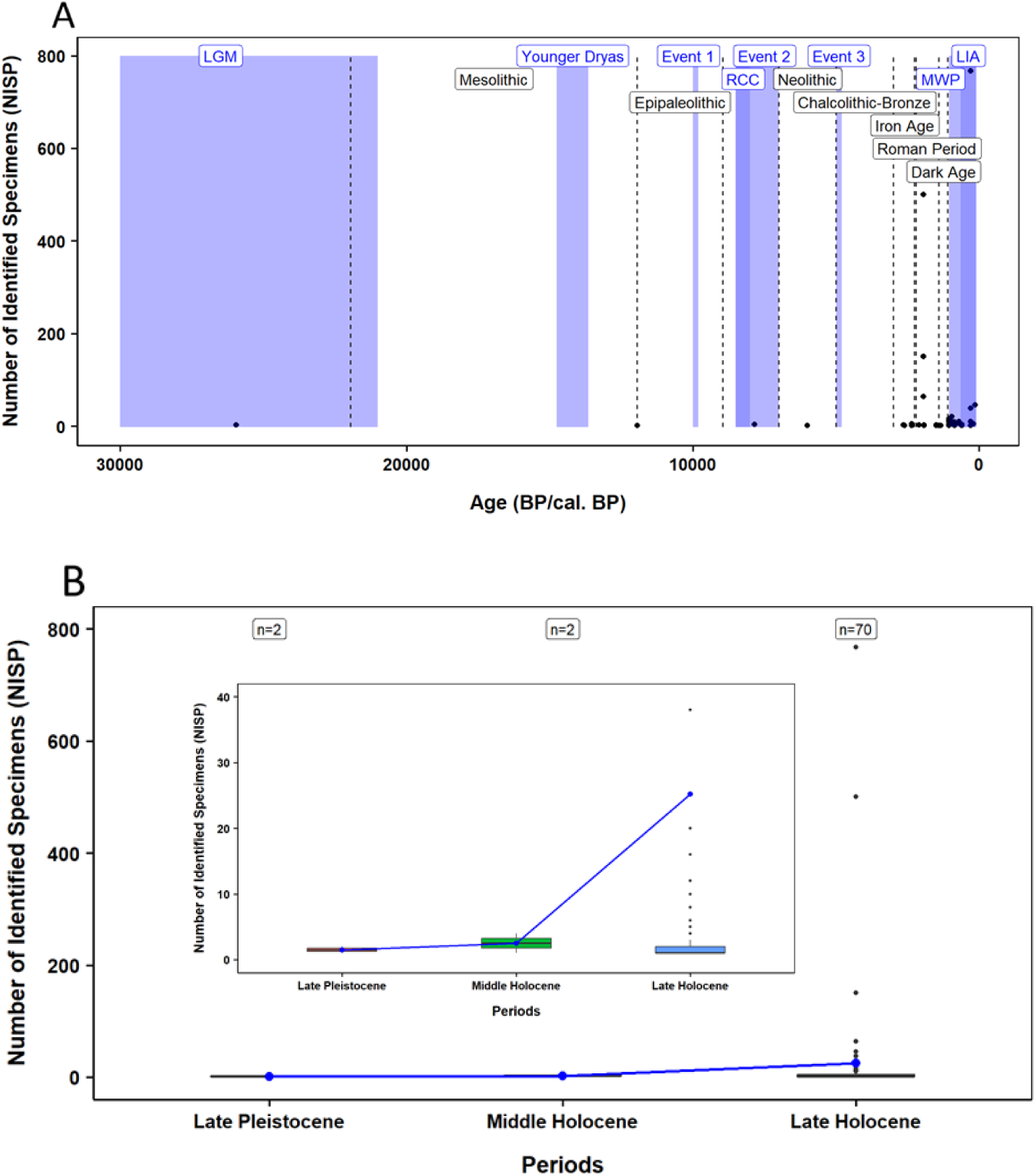
Distribution of data for NISP on the Atlantic bluefin tuna in the Western Mediterranean. (A) Temporal distribution of records, including major climatic events (purple rectangles) and anthropogenic events (black dashed lines); (B) Boxplots of NISP data for each geological period. Blue dots represent mean values, and bars indicate standard deviation. Letters denote statistically significant differences between periods. The number of data samples (n) for each boxplot is shown above. The plot inside the second figure is a zoom-in of the data, for better visualisation of the increase in NISP values towards the Late Holocene (no outliers were excluded).

**Figure S3.2-.**
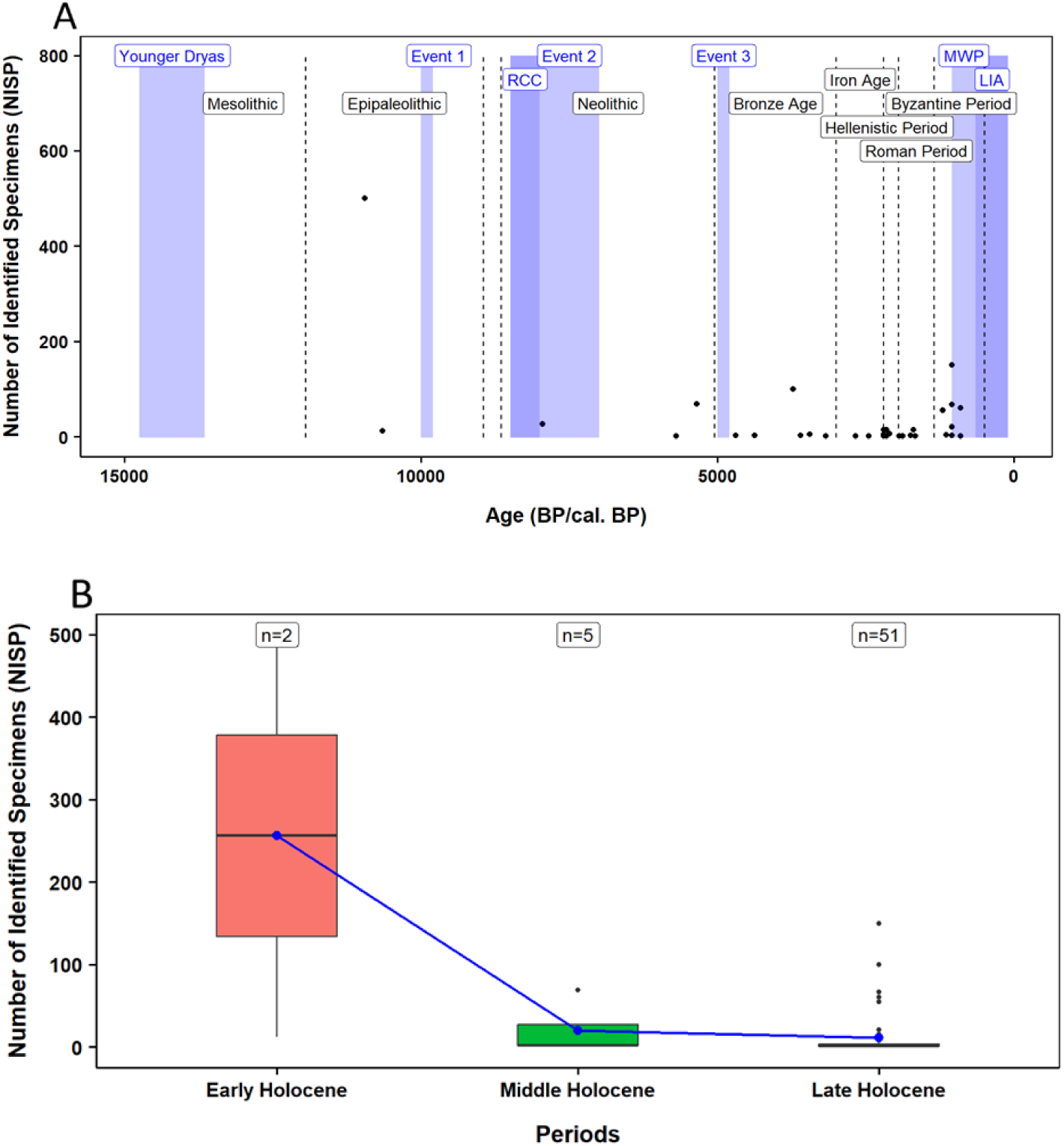
Distribution of data for NISP on the Atlantic bluefin tuna in the Eastern Mediterranean. (A) Temporal distribution of records, including major climatic events (purple rectangles) and anthropogenic events (black dashed lines); (B) Boxplots of NISP data for each geological period. Blue dots represent mean values, and bars indicate standard deviation. Letters denote statistically significant differences between periods. The number of data samples (n) for each boxplot is shown above.

**Figure S3.3-.**
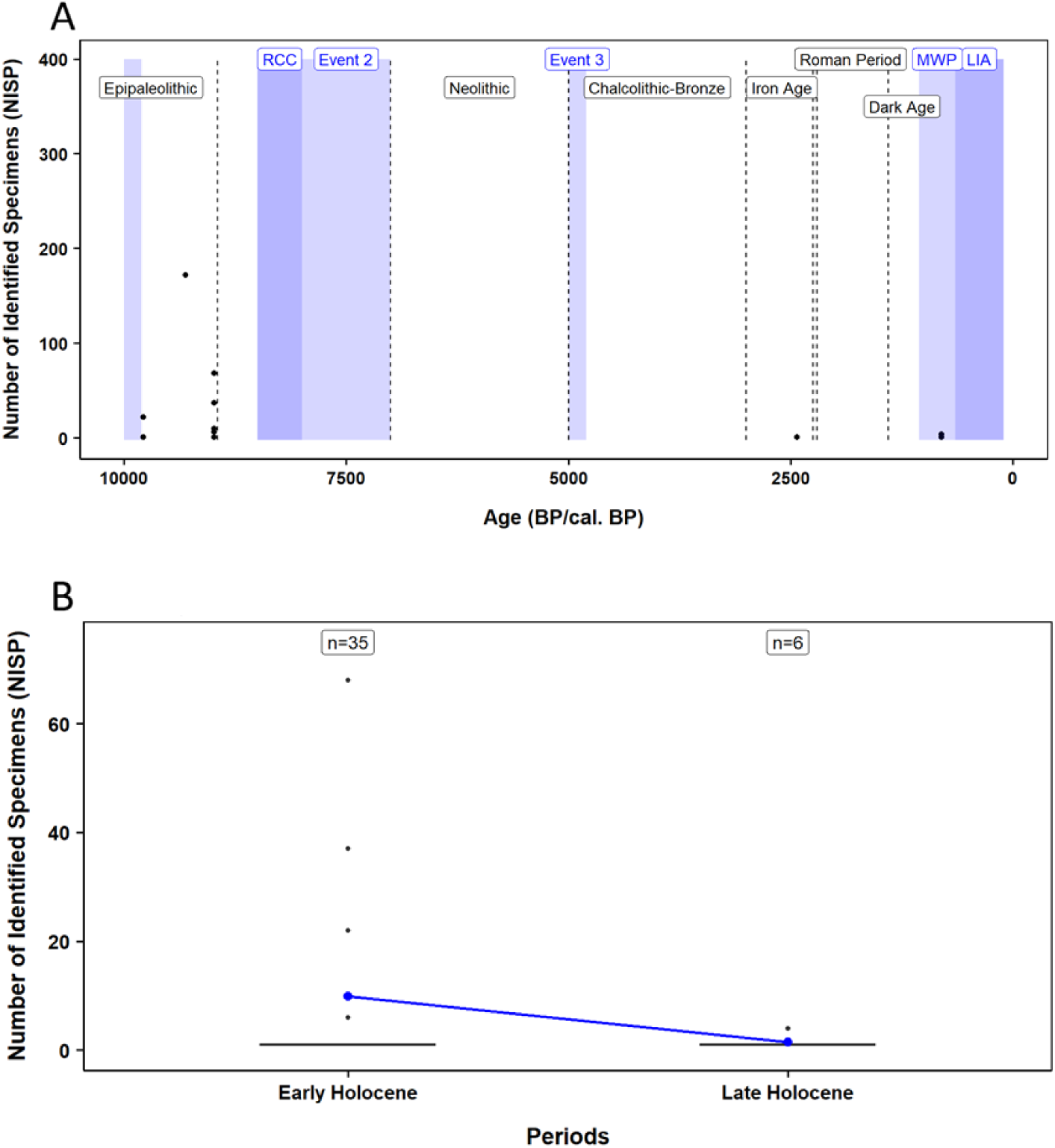
Distribution of data for NISP on the gilthead sea bream in the Western Mediterranean. (A) Temporal distribution of records, including major climatic events (purple rectangles) and anthropogenic events (black dashed lines); (B) Boxplots of NISP data for each geological period. Blue dots represent mean values, and bars indicate standard deviation. Letters denote statistically significant differences between periods. The number of data samples (n) for each boxplot is shown above.

**Figure S3.4-.**
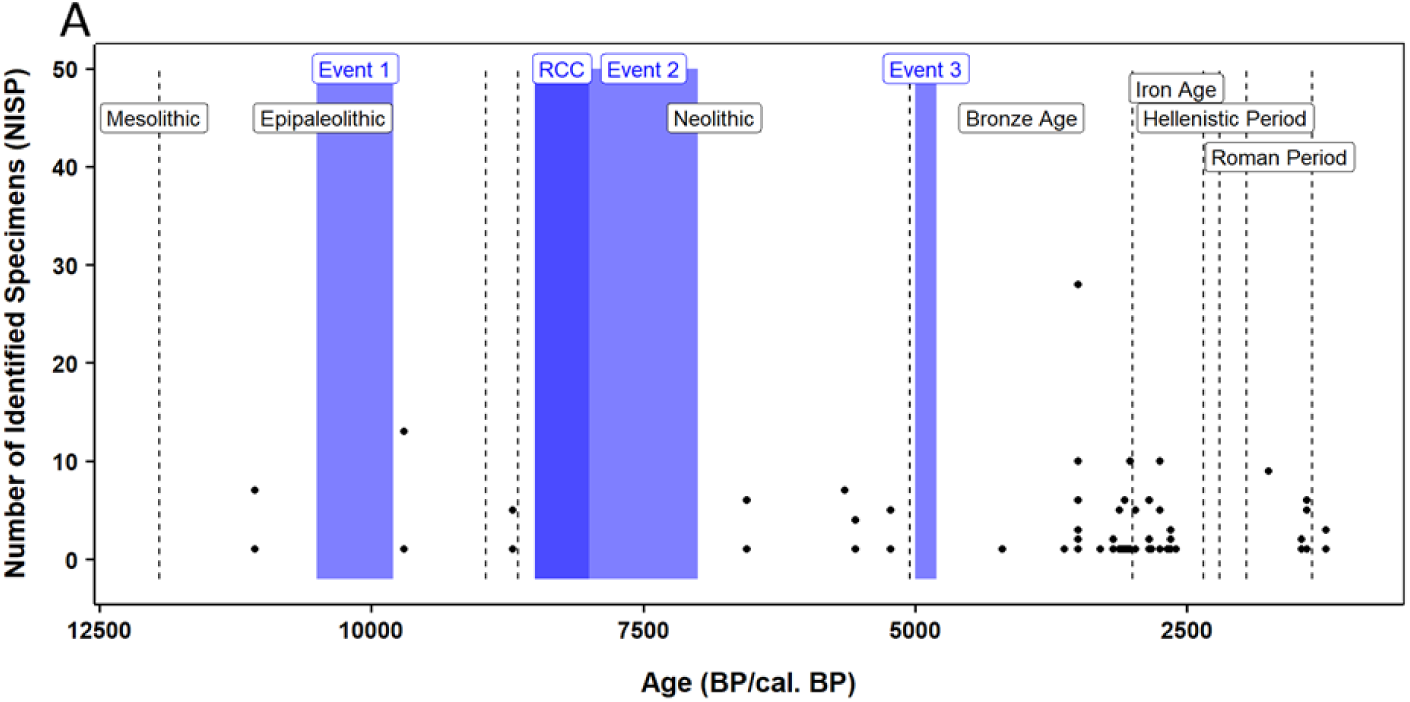
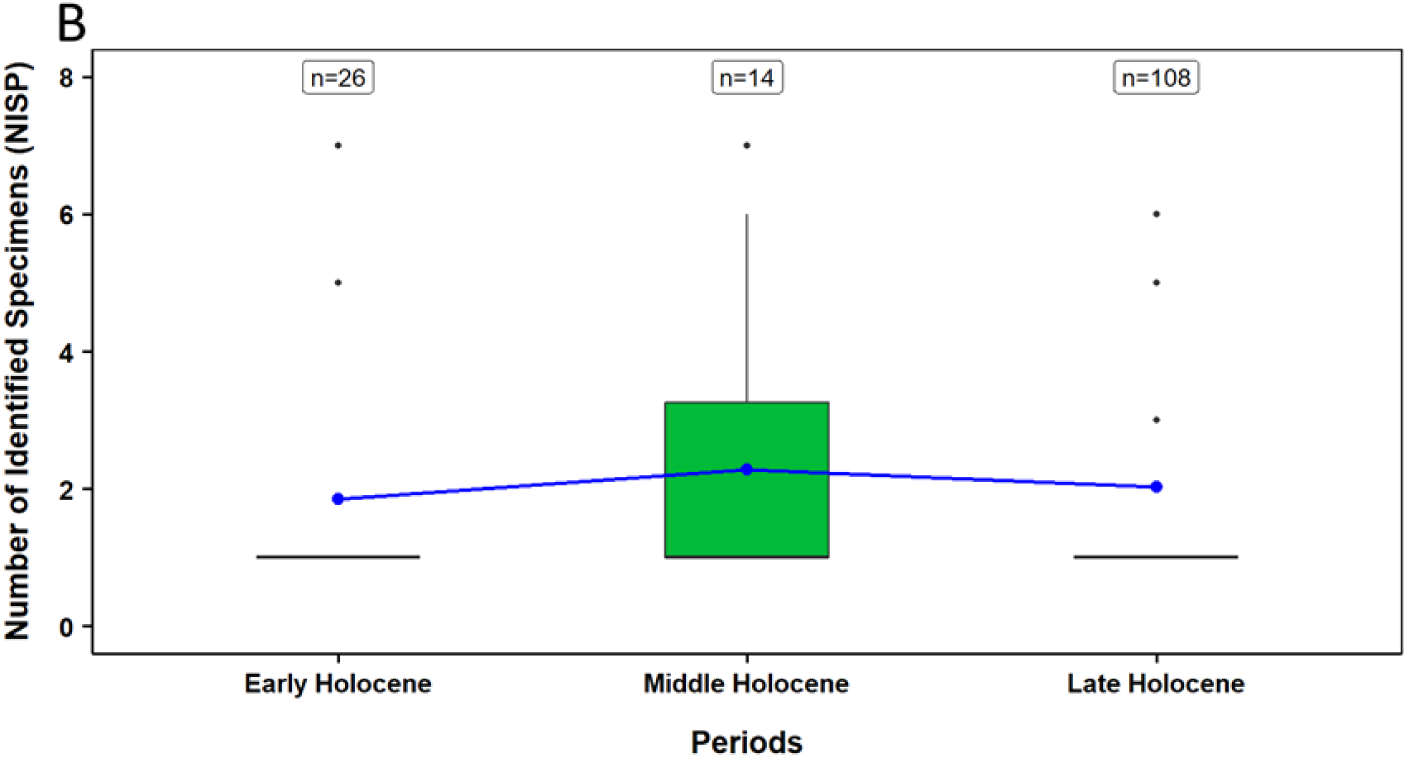
Distribution of data for NISP on the gilthead sea bream in the Eastern Mediterranean. (A) Temporal distribution of records, including major climatic events (purple rectangles) and anthropogenic events (black dashed lines); (B) Boxplots of NISP data for each geological period. Blue dots represent mean values, and bars indicate standard deviation. Letters denote statistically significant differences between periods. The number of data samples (n) for each boxplot is shown above.

**Figure S3.5-.**
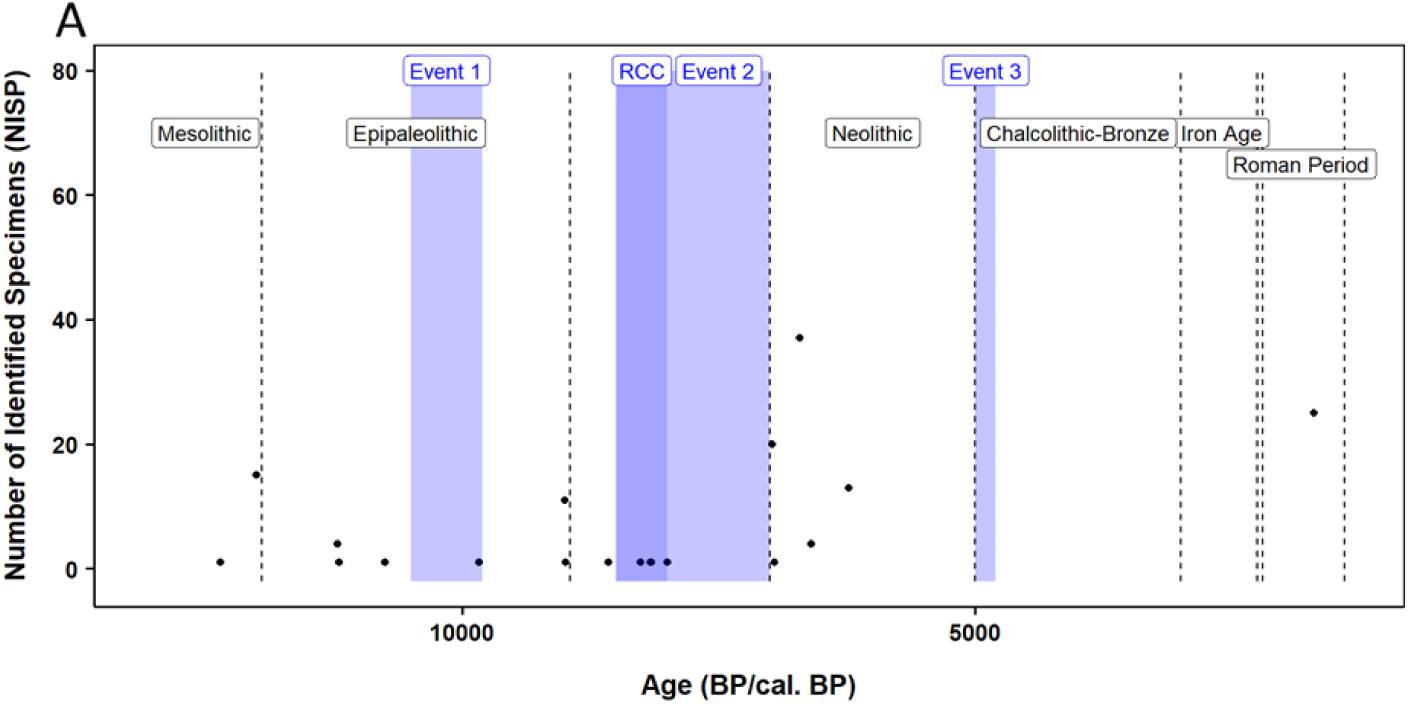
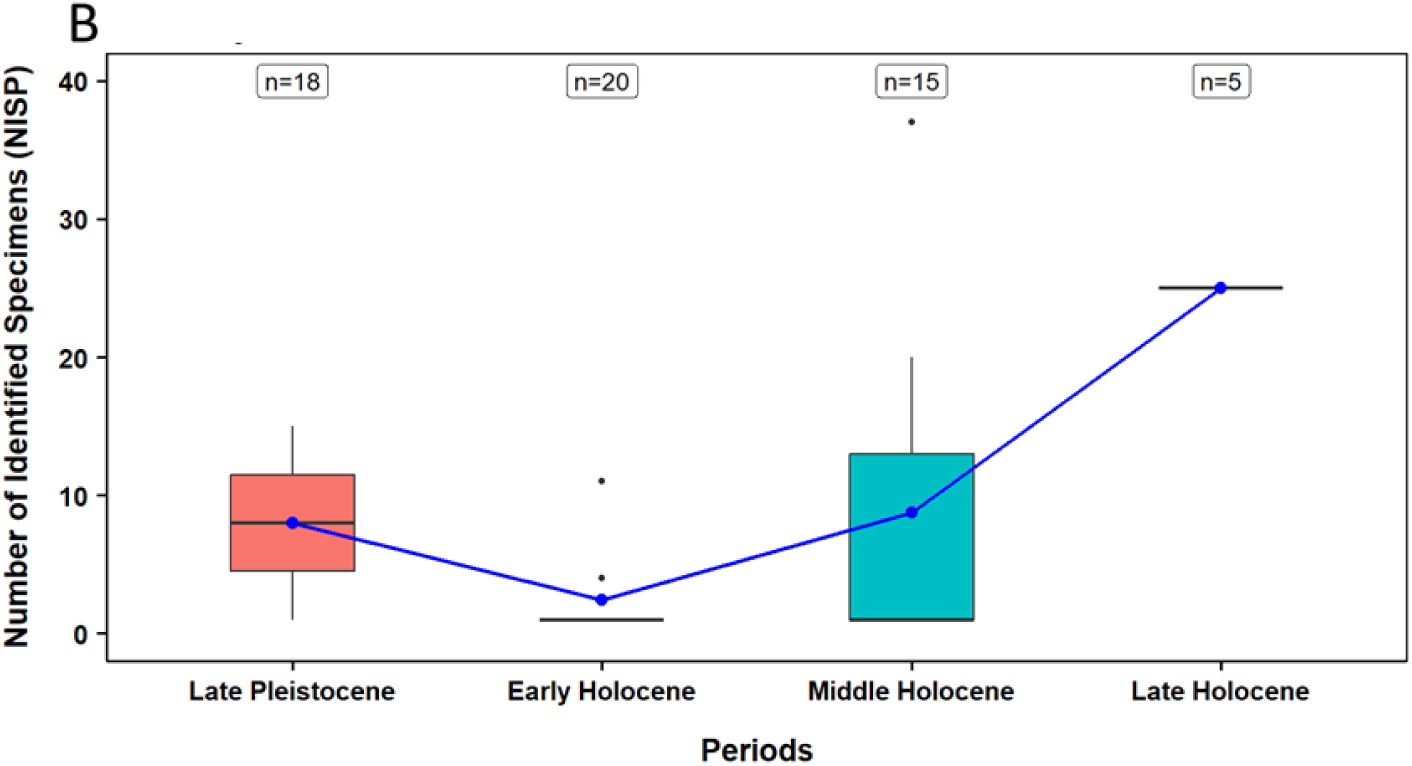
Distribution of data for NISP on the turbinate monodont in the Western Mediterranean. (A) Temporal distribution of records, including major climatic events (purple rectangles) and anthropogenic events (black dashed lines); (B) Boxplots of NISP data for each geological period. Blue dots represent mean values, and bars indicate standard deviation. Letters denote statistically significant differences between periods. The number of data samples (n) for each boxplot is shown above.

**Figure S3.6-.**
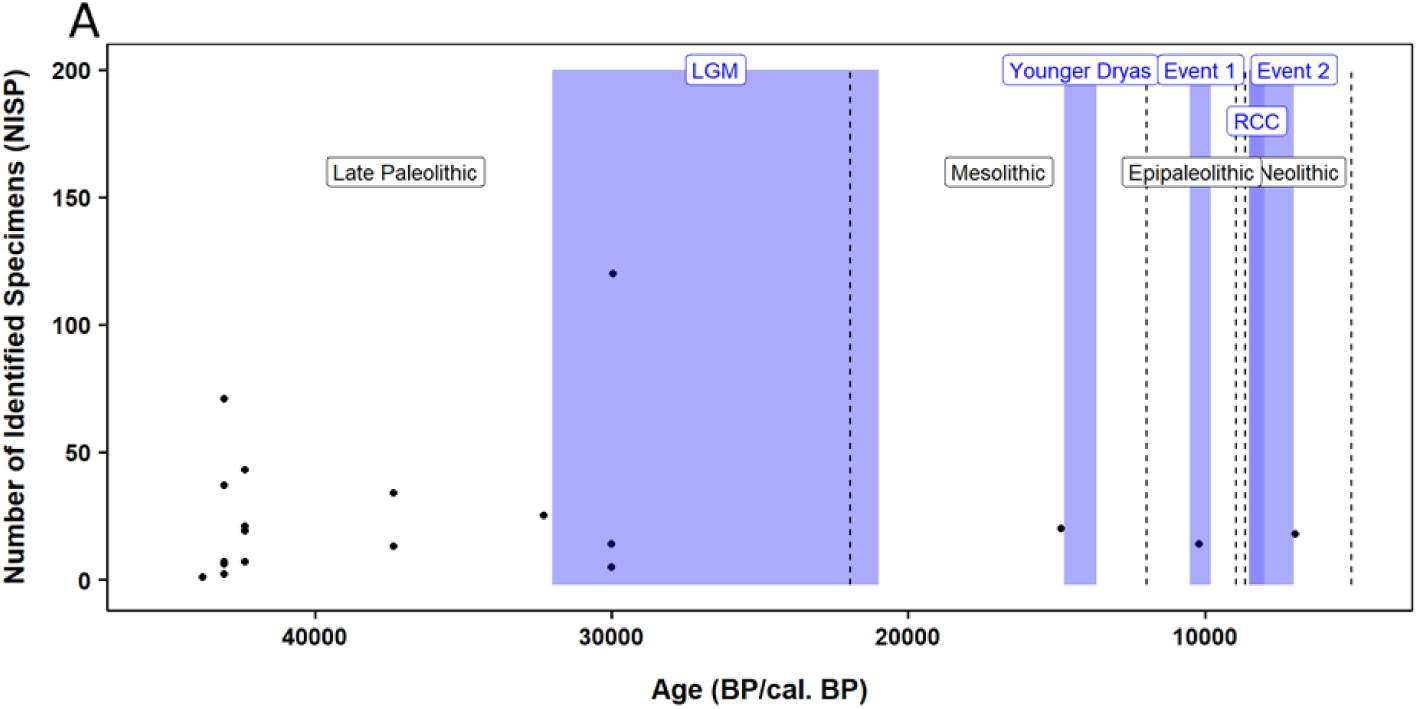
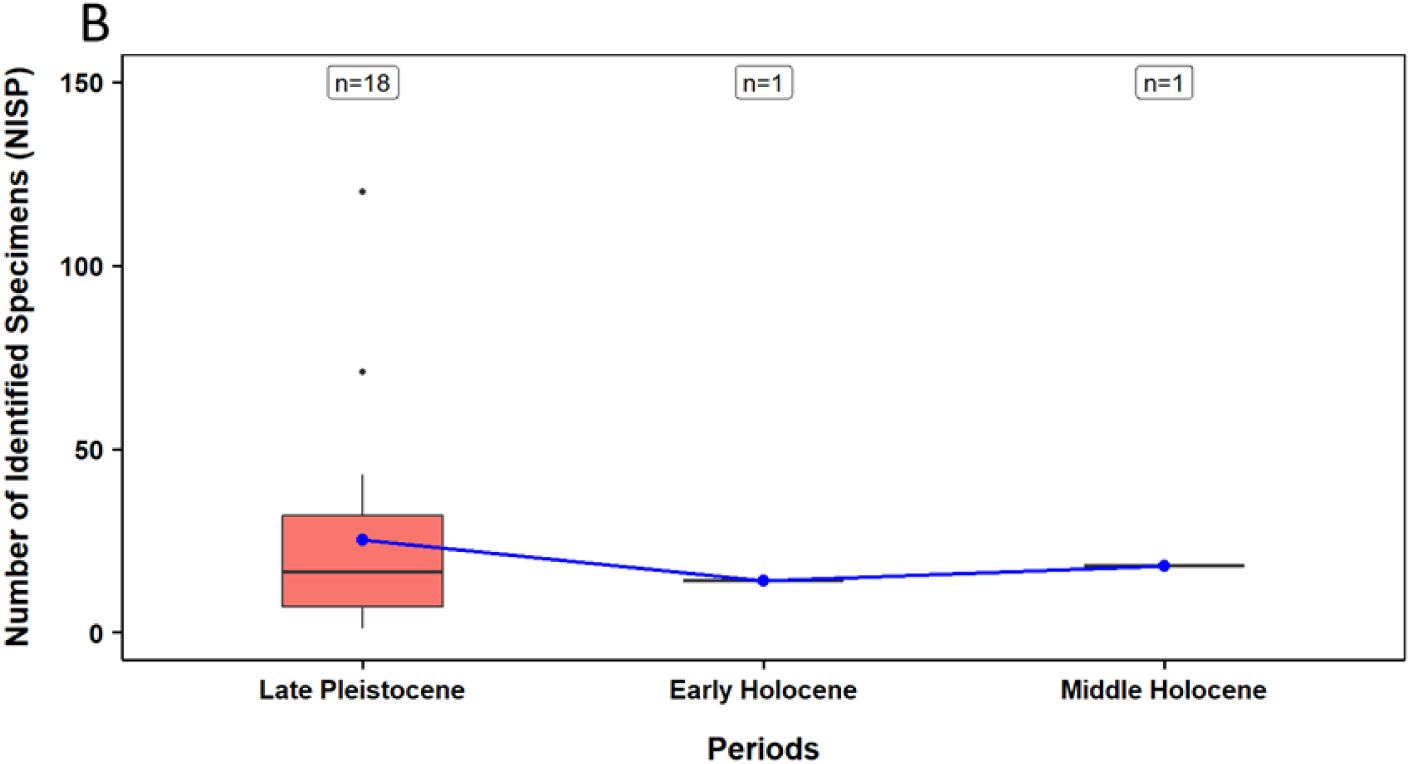
Distribution of data for NISP on the turbinate monodont in the Eastern Mediterranean. (A) Temporal distribution of records, including major climatic events (purple rectangles) and anthropogenic events (black dashed lines); (B) Boxplots of NISP data for each geological period. Blue dots represent mean values, and bars indicate standard deviation. Letters denote statistically significant differences between periods. The number of data samples (n) for each boxplot is shown above.

**Figure S3.7-.**
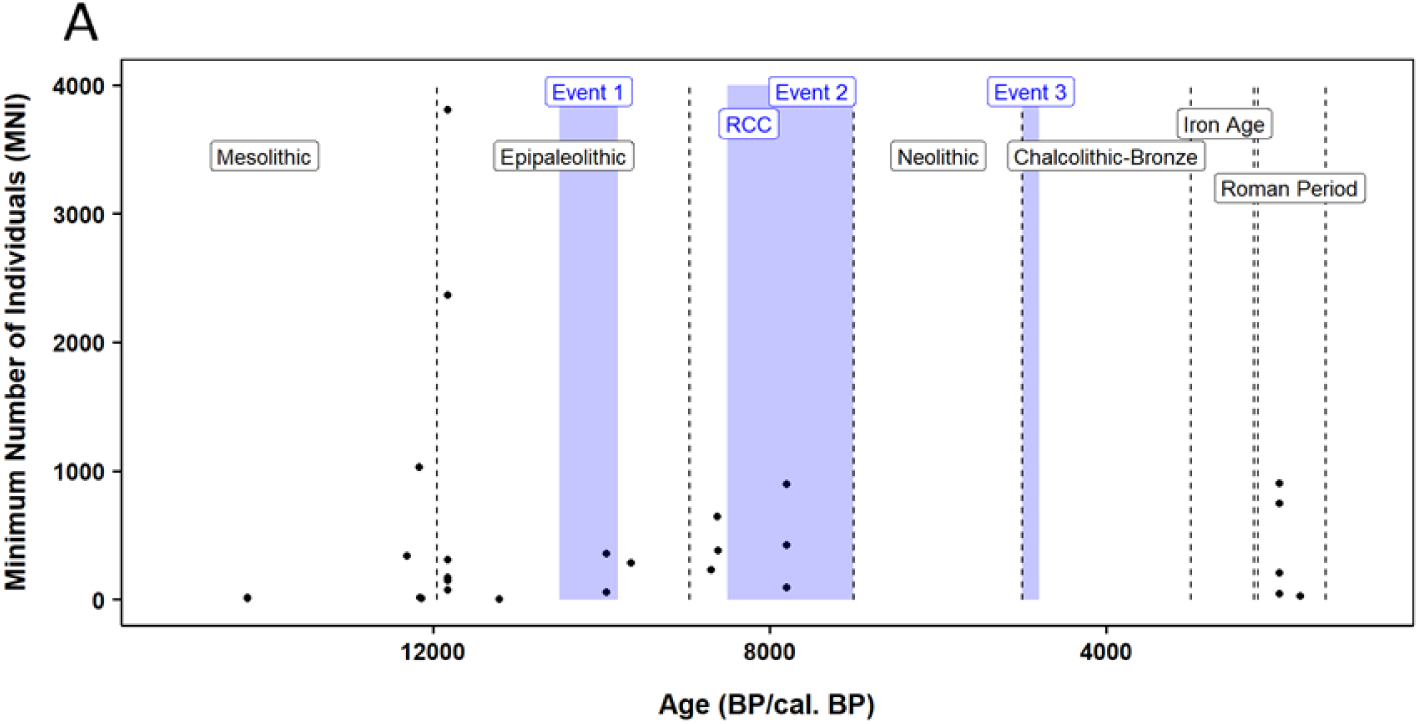
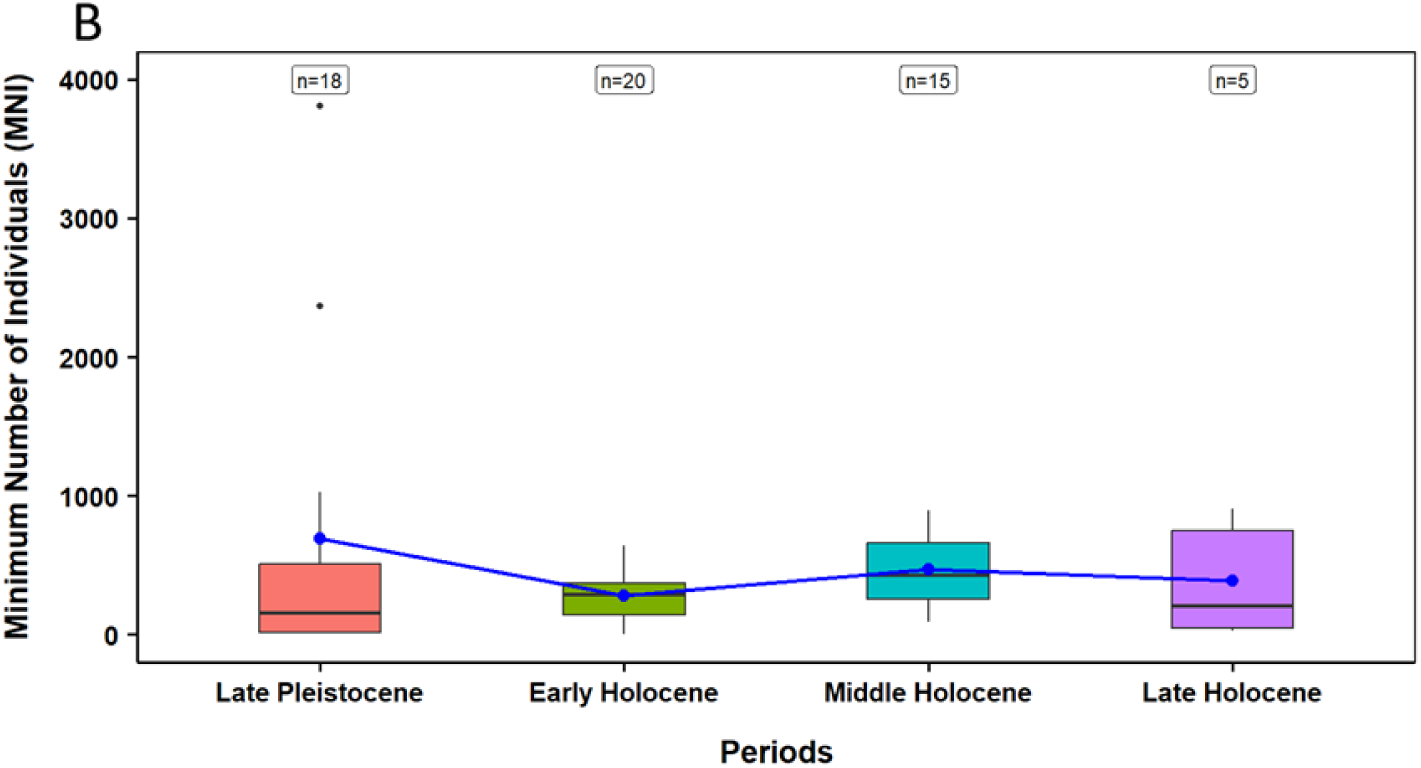
Distribution of data for MNI on the turbinate monodont in the Western Mediterranean. (A) Temporal distribution of records, including major climatic events (purple rectangles) and anthropogenic events (black dashed lines); (B) Boxplots of MNI data for each geological period. Blue dots represent mean values, and bars indicate standard deviation. Letters denote statistically significant differences between periods. The number of data samples (n) for each boxplot is shown above.

**Figure S3.8-.**
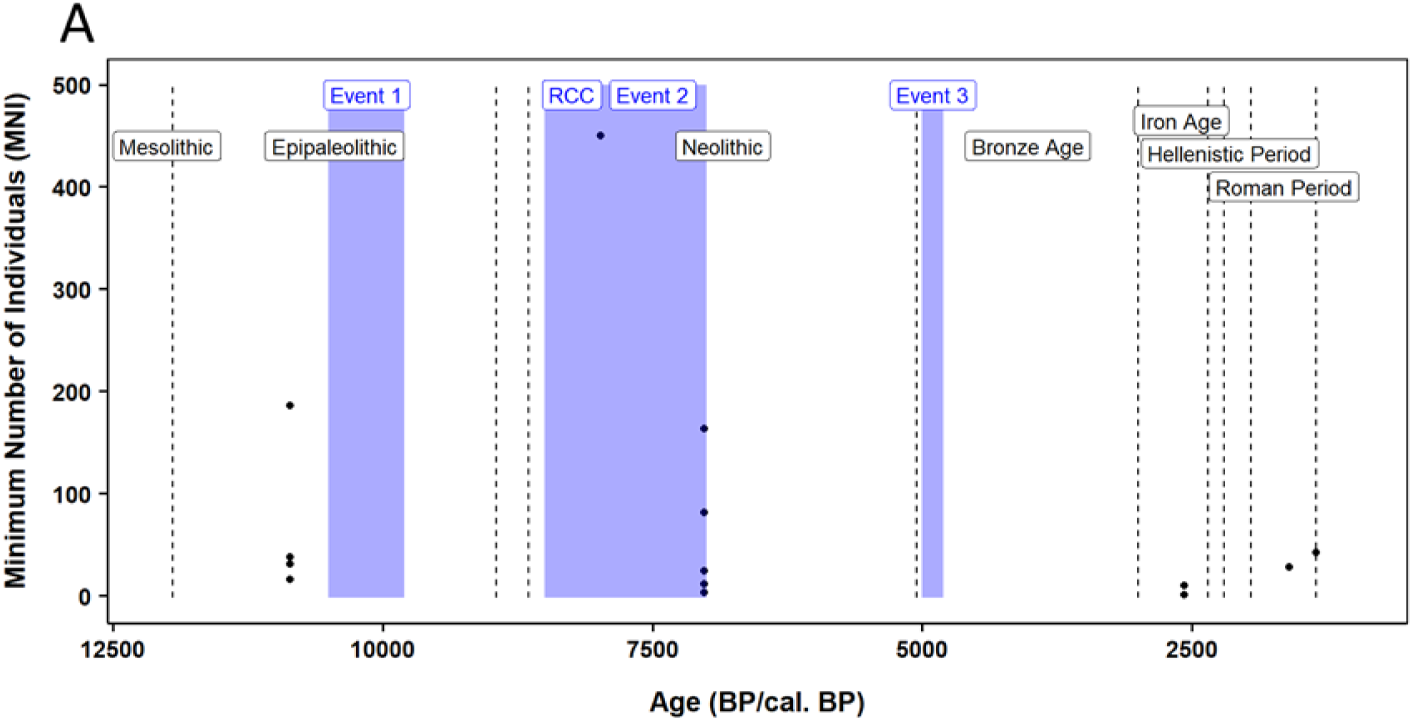
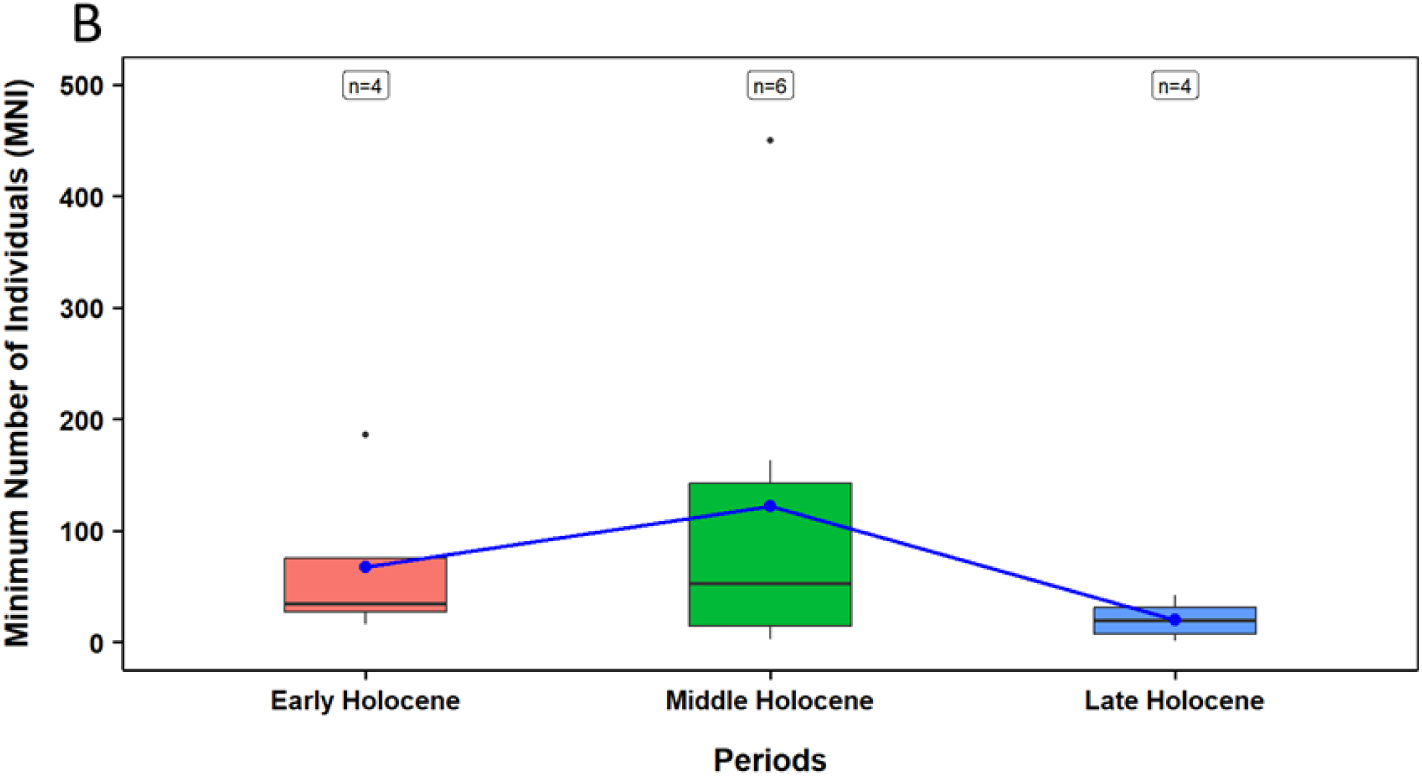
Distribution of data for MNI on the turbinate monodont in the Eastern Mediterranean. (A) Temporal distribution of records, including major climatic events (purple rectangles) and anthropogenic events (black dashed lines); (B) Boxplots of MNI data for each geological period. Blue dots represent mean values, and bars indicate standard deviation. Letters denote statistically significant differences between periods. The number of data samples (n) for each boxplot is shown above.

### S4. Catch records

**Figure S4-.**
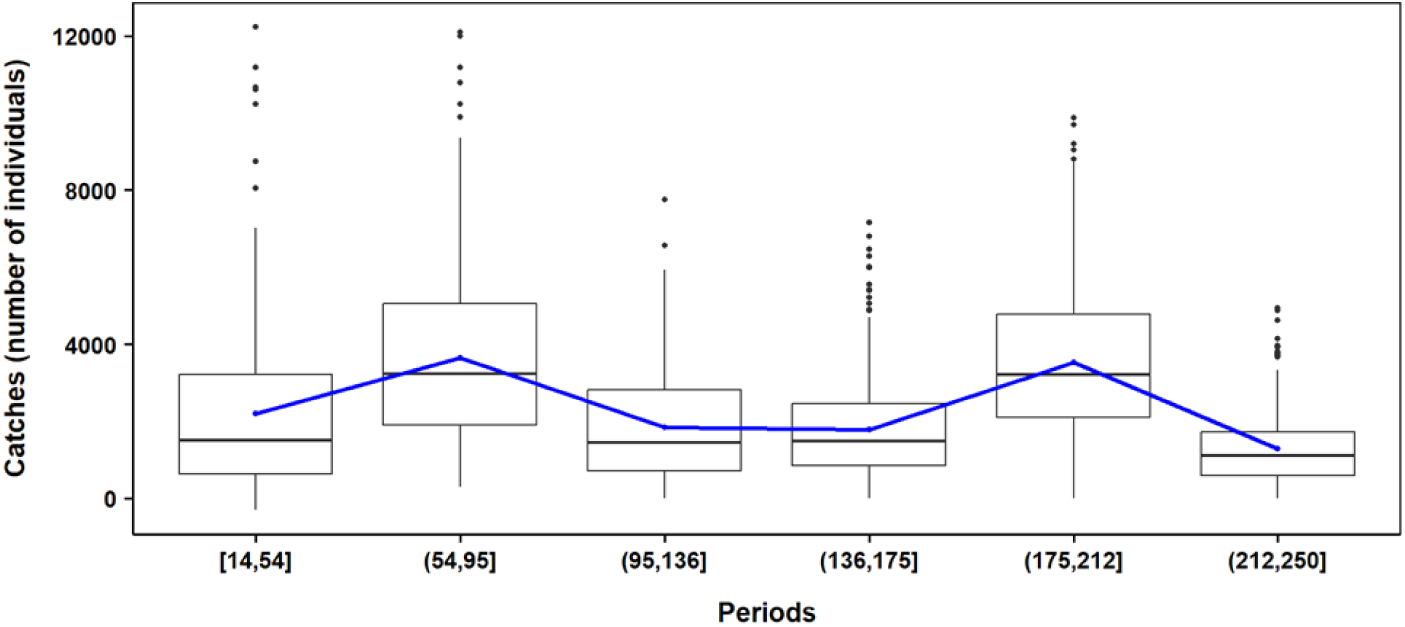
Catch records of the Atlantic bluefin tuna are divided into six periods, using the Jenks Natural Breaks Method, between 250–14 years BP. The blue dots, connected by the blue line, are the mean values for each period. The number of samples (n) included in each box plot is indicated above. The black dots are outliers.

**Table S4-.** Pairwise comparisons between intervals (years BP) for the Atlantic bluefin tuna catch numbers. Numbers in bold represent the periods that differ significantly from each other.

|  | [14,54] | (54, 95] | (95,136] | (136,175] | (175,212] |
| --- | --- | --- | --- | --- | --- |
| (54, 95] | $< 2e^{-16}$ | | | | |
| (95,136] | 1 | $< 2e^{-16}$ | | | |
| (136,175] | 1 | $< 2e^{-16}$ | 1 | | |
| (175,212] | $< 2e^{-16}$ | 1 | $< 2e^{-16}$ | $< 2e^{-16}$ | |
| (212,250] | $6e^{-04}$ | $< 2e^{-16}$ | $3e^{-04}$ | $1.1e^{-05}$ | $< 2e^{-16}$ |

### S5. Body size

**Figure S5.1-.**
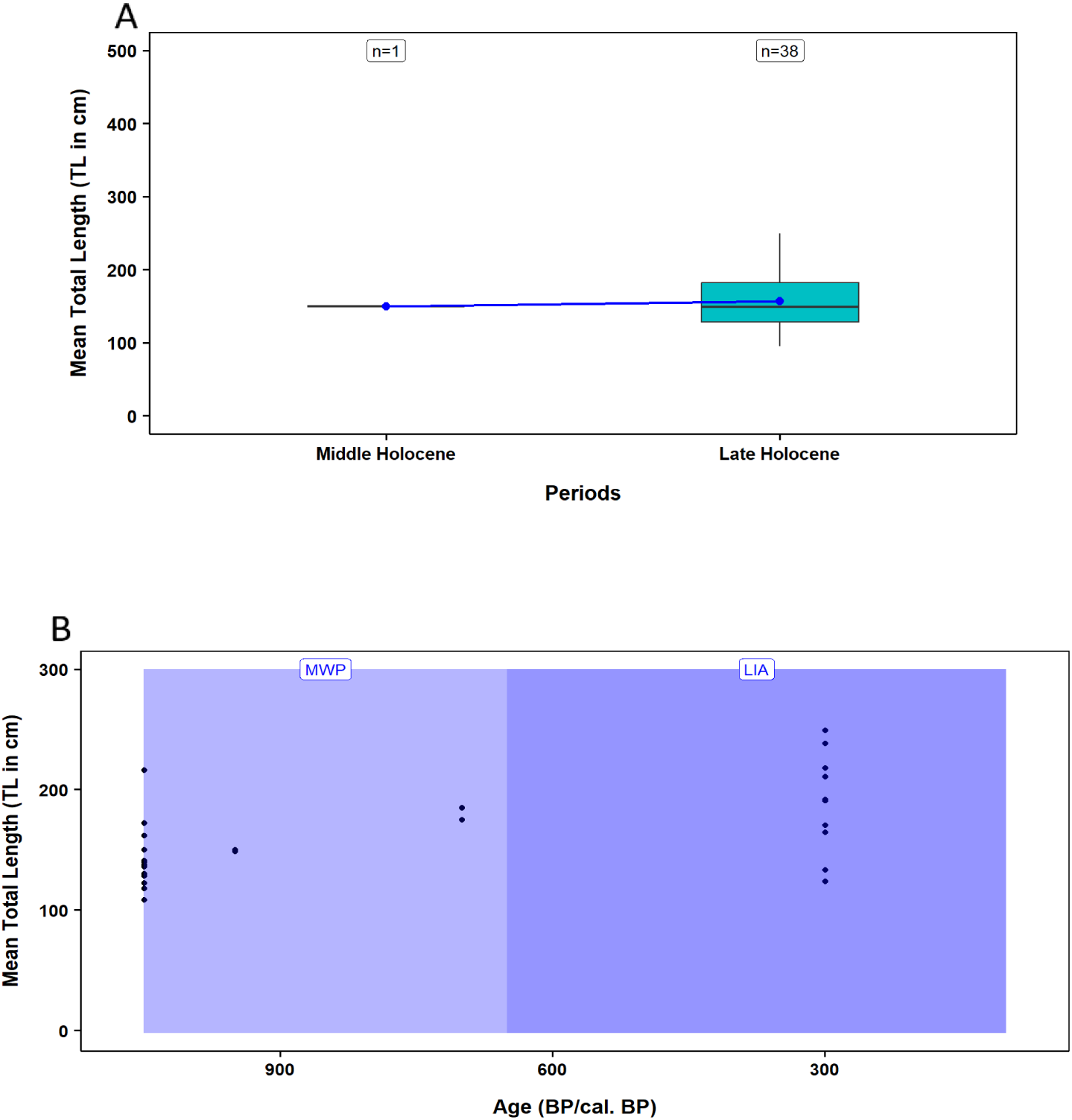
Distribution of data for TL on the Atlantic bluefin tuna in the Western Mediterranean. (A) Boxplots of TL data for each geological period. Blue dots represent mean values, and bars indicate standard deviation. Letters denote statistically significant differences between periods. The number of data samples (n) for each boxplot is shown above. (B) Total length differences between the MWP (Medieval Warm Period) and LIA (Little Ice Age).

**Figure S5.2-.**
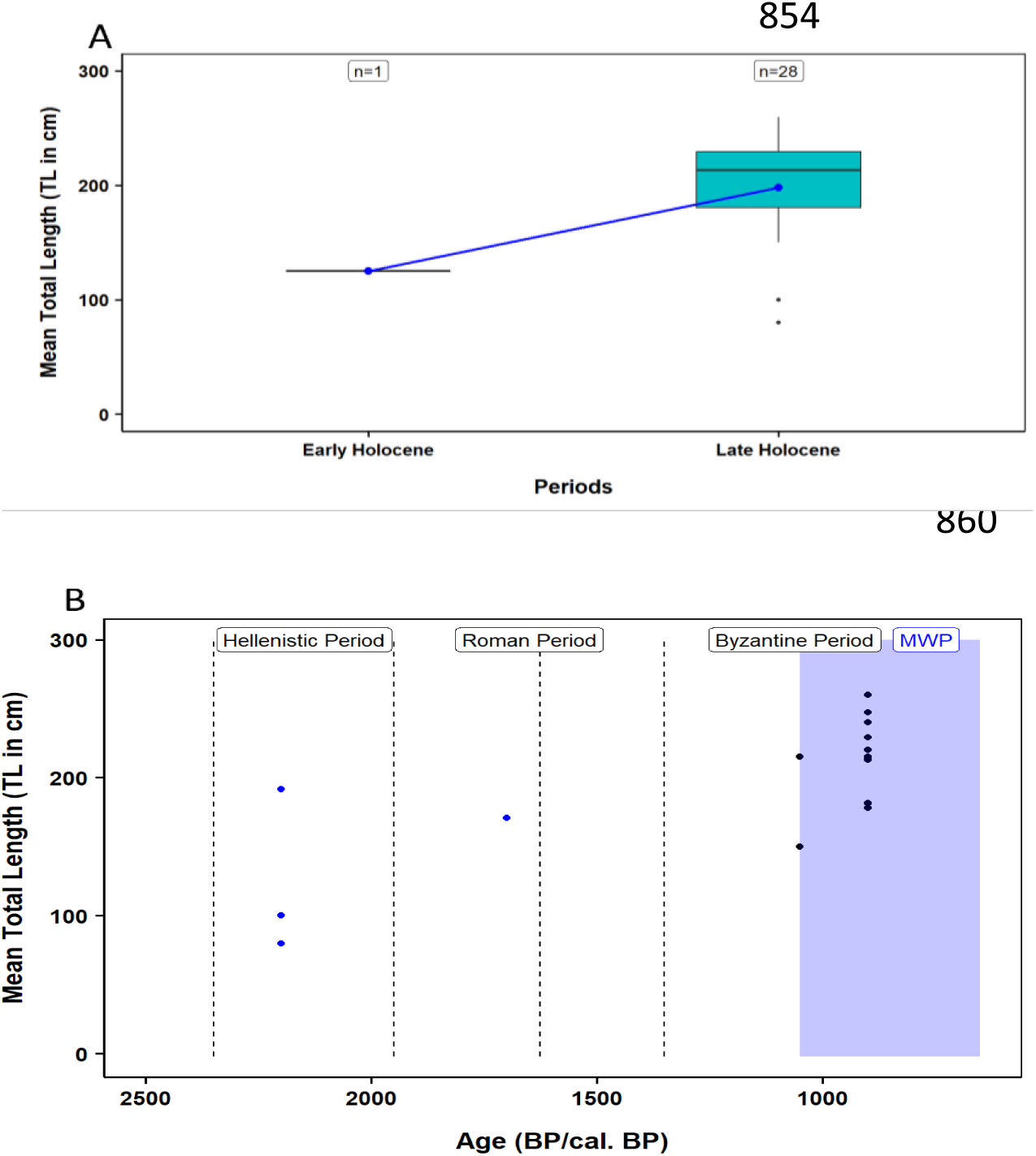
Distribution of data for TL on the Atlantic bluefin tuna in the Eastern Mediterranean. (A) Boxplots of TL data for each geological period. Blue dots represent mean values, and bars indicate standard deviation. Letters denote statistically significant differences between periods. The number of data samples (n) for each boxplot is shown above. (B) differences between the before and during the MWP (Medieval Warm Period).

**Figure S5.3-.**
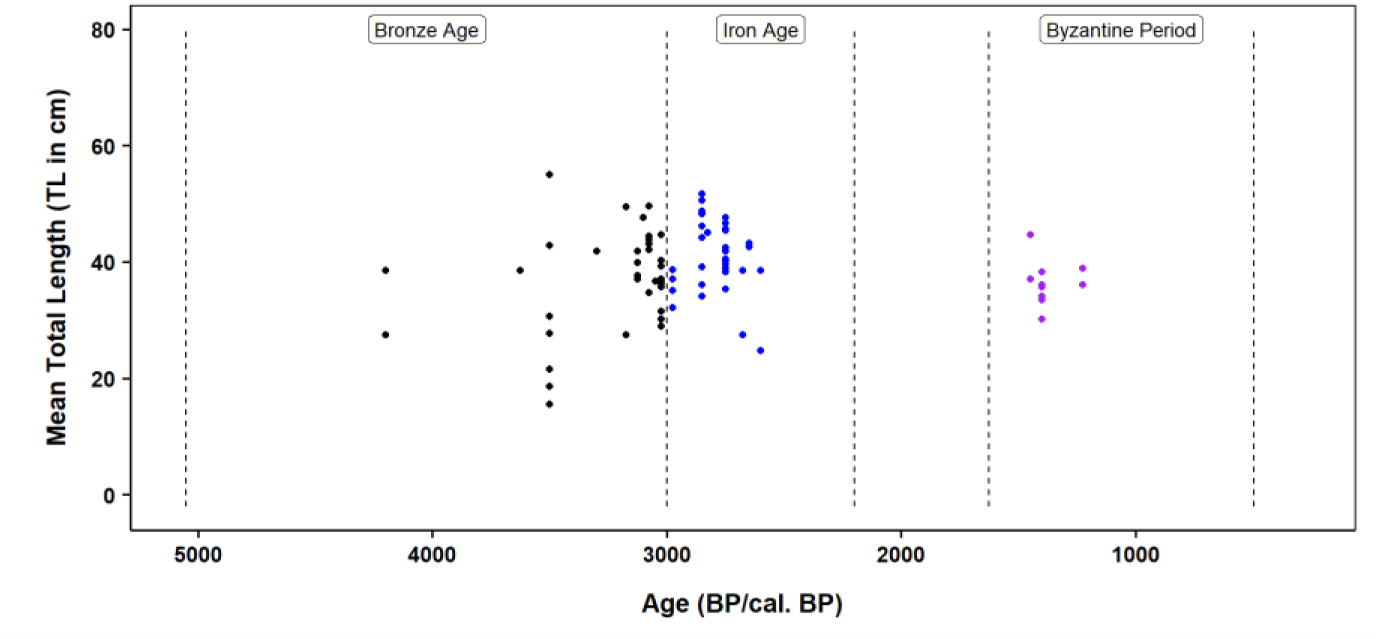
Distribution of data for the TL on the Gilthead sea bream in the Eastern Mediterranean. Differences between the Bronze and Iron Ages and the Byzantine period.

**Figure S5.4-.**
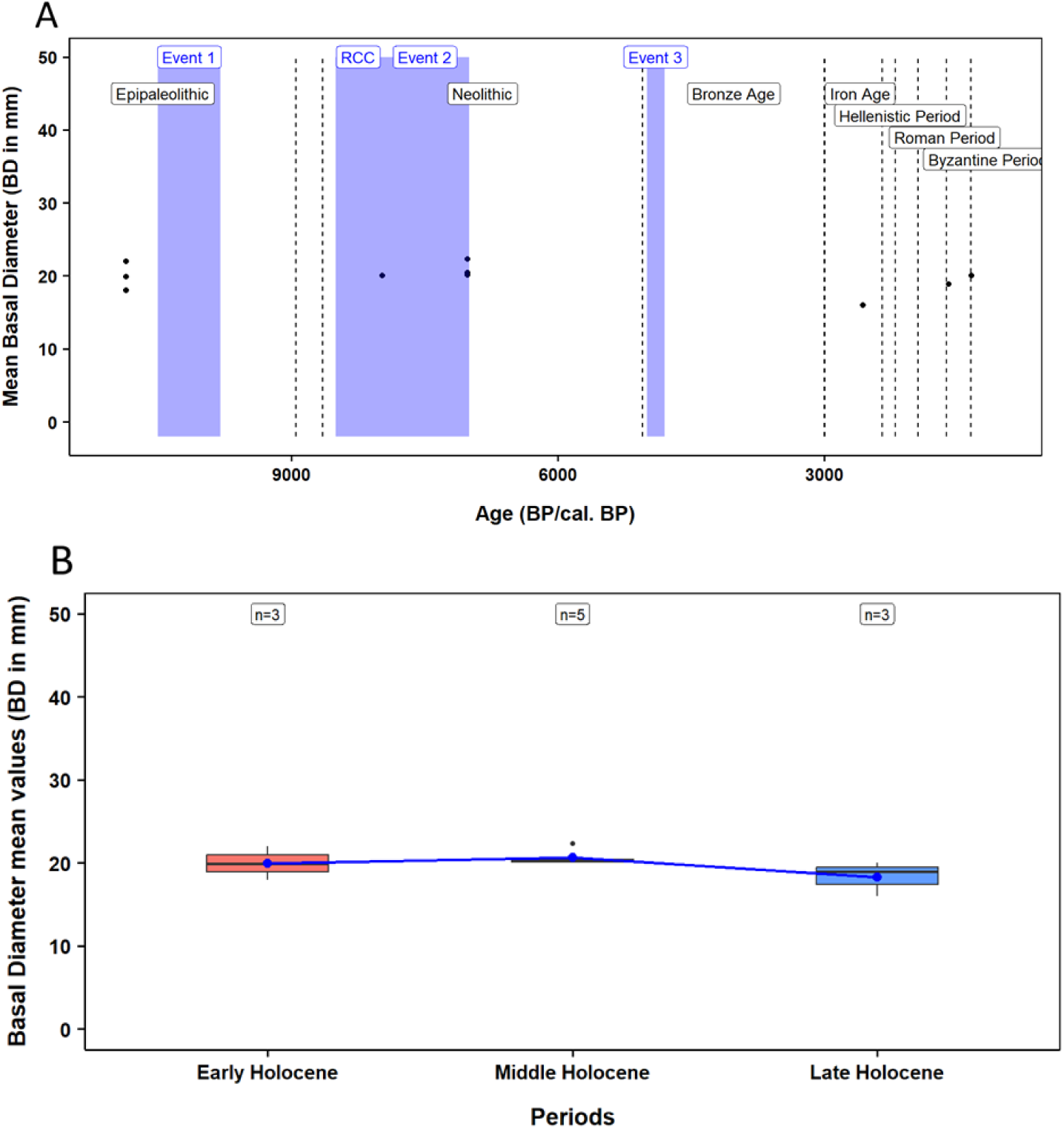
Distribution of data for mean BD on the turbinate monodont in the Eastern Mediterranean. (A) Temporal distribution of records, including major climatic events (purple rectangles) and anthropogenic events (black dashed lines); (B) Boxplots of BD data for each geological period. Blue dots represent mean values, and bars indicate standard deviation. Letters denote statistically significant differences between periods. The number of data samples (n) for each boxplot is shown above.

**Figure S5.5-.**
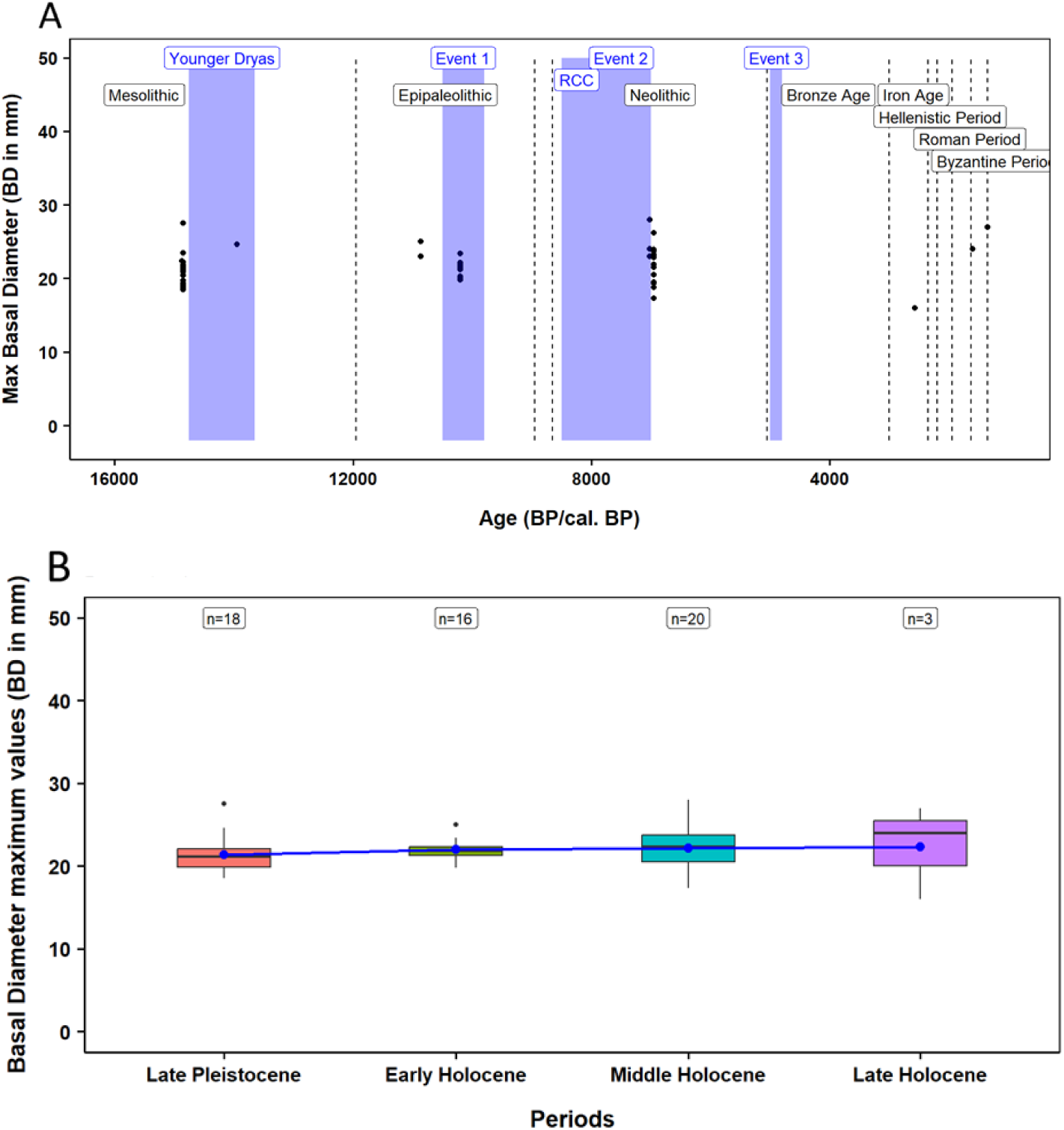
Distribution of data for maximum values of BD on the turbinate monodont in the Eastern Mediterranean. (A) Temporal distribution of records, including major climatic events (purple rectangles) and anthropogenic events (black dashed lines); (B) Boxplots of BD data for each geological period. Blue dots represent mean values, and bars indicate standard deviation. Letters denote statistically significant differences between periods. The number of data samples (n) for each boxplot is shown above.

### S6. Sampling site and references

**Table S6-.**
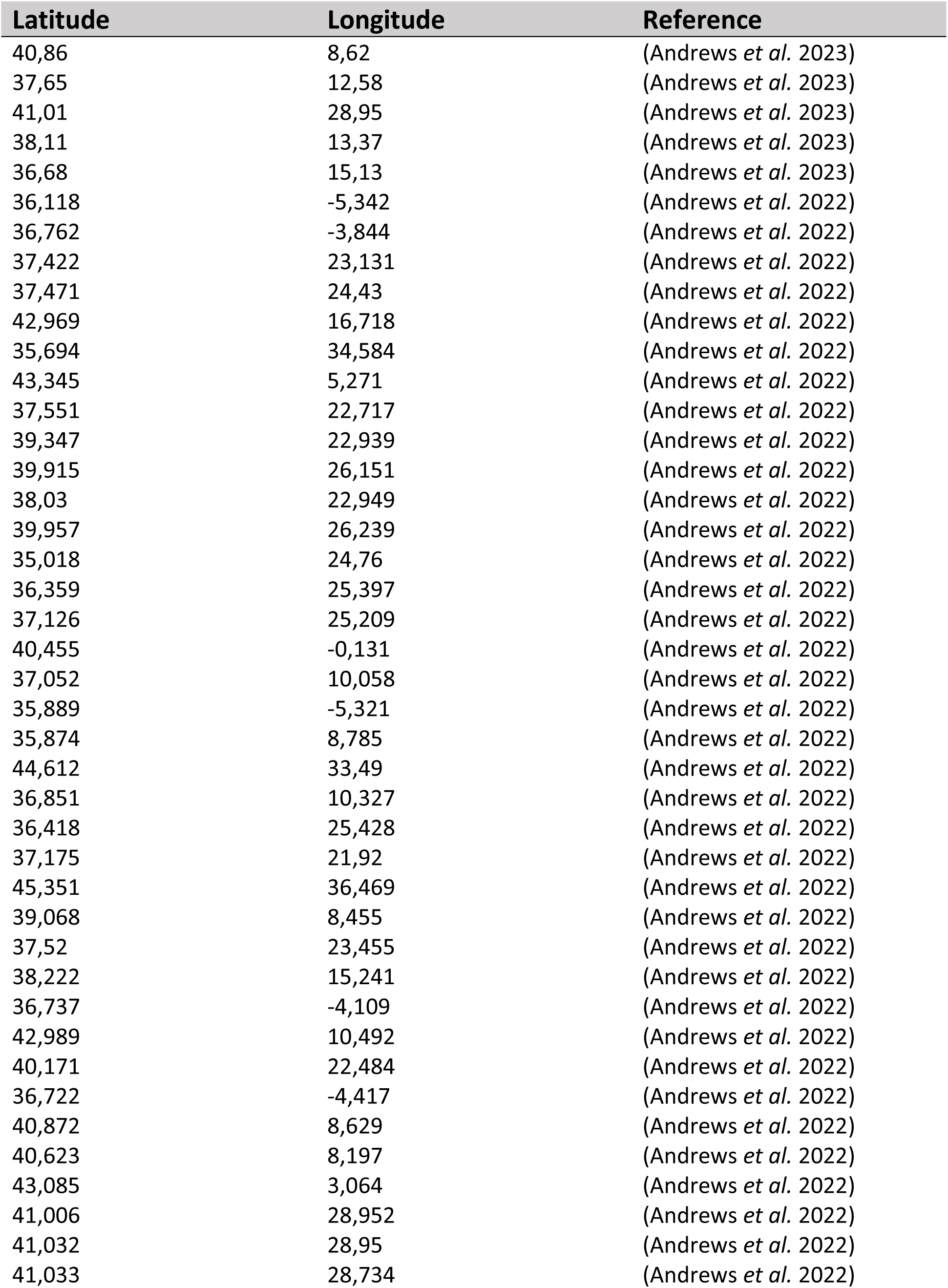
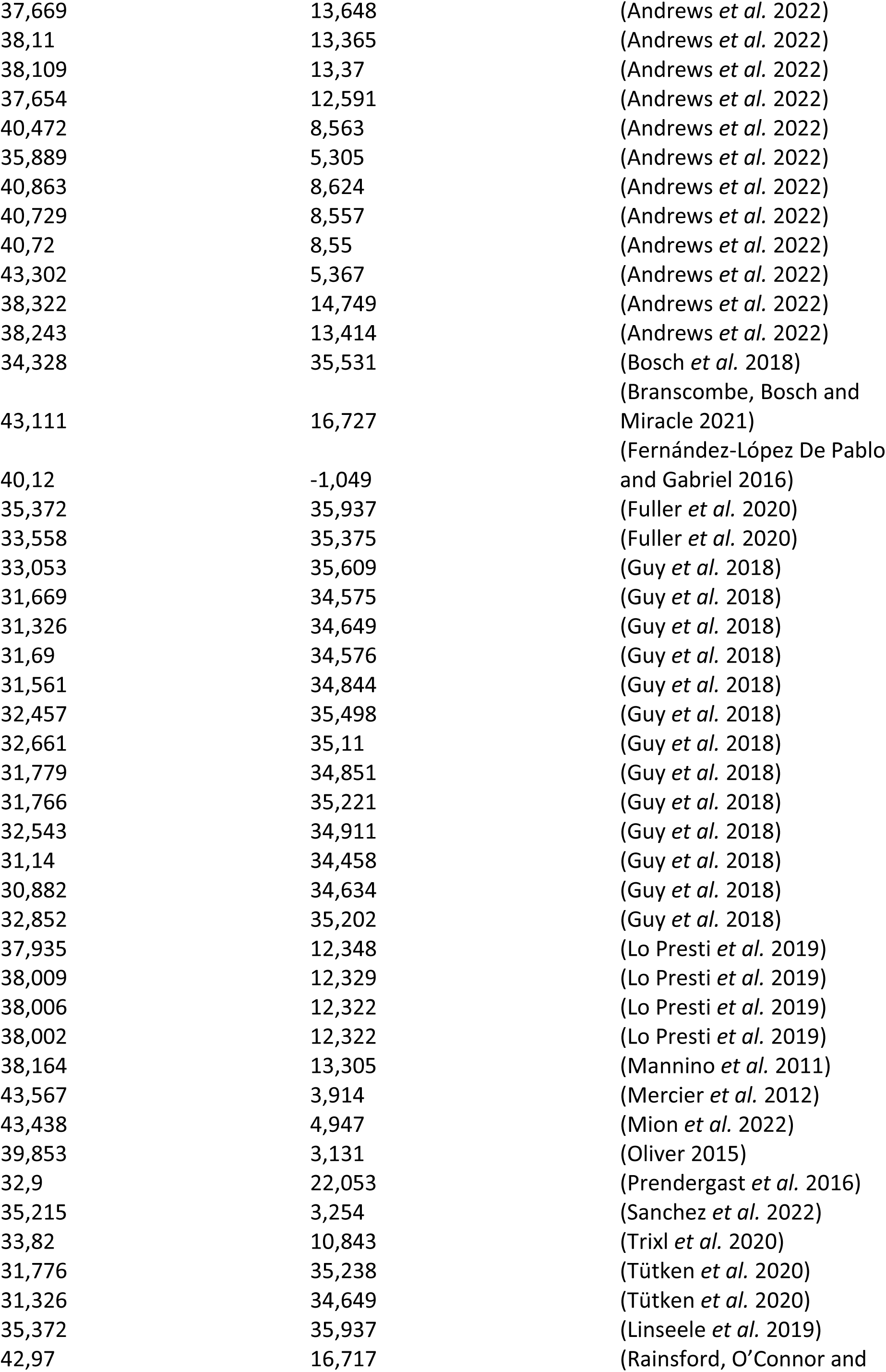
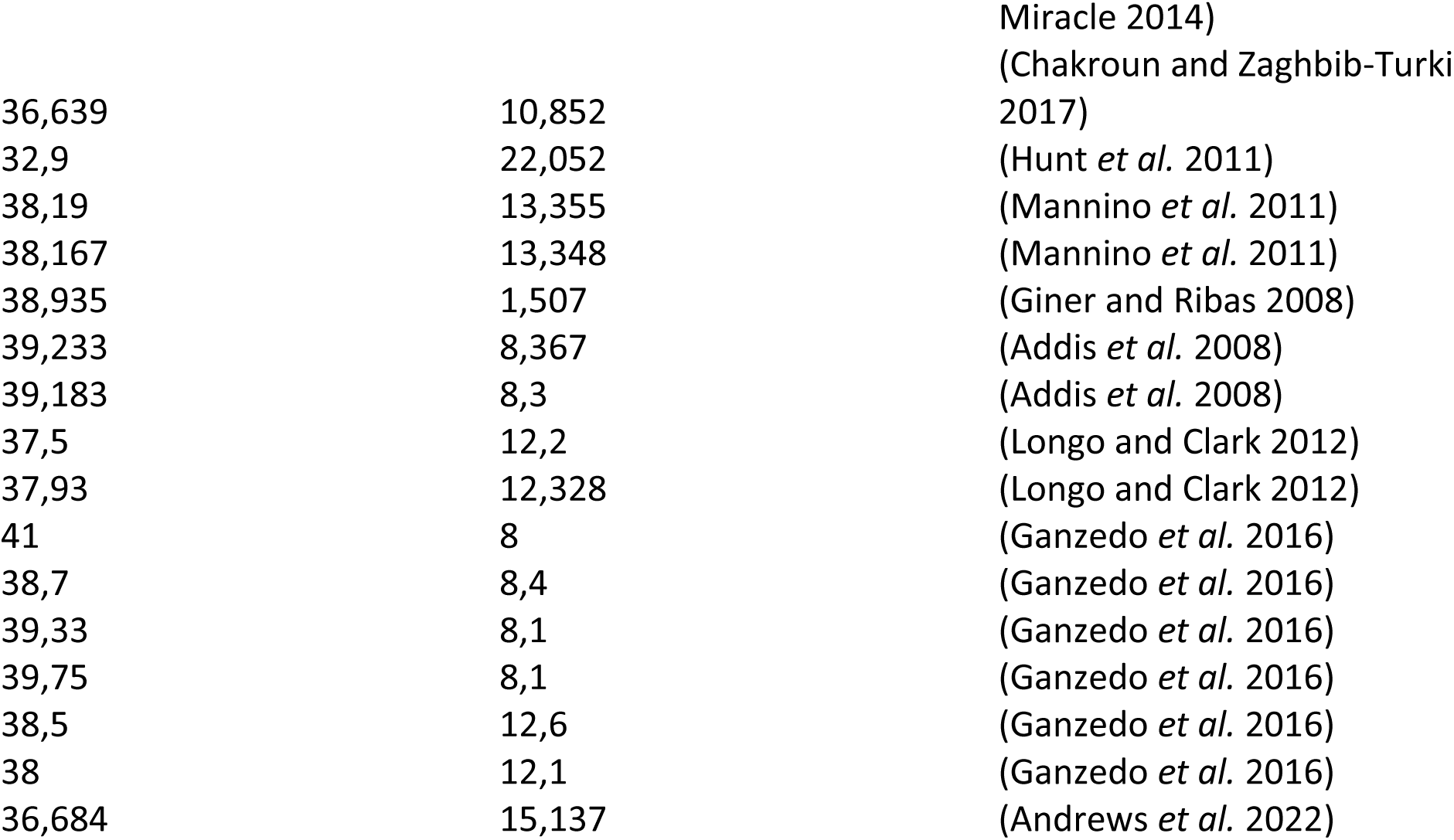
Table with information on the coordinates of each sampling site, with corresponding variable from which data were extracted, and the literary reference.

### S7. Species’ number of samples per variable

**Table S7-.** Information on the number of samples (n) per variable for each species. NISP – Number of Identified Specimens; MNI – Minimum Number of Species; TL – Total Length; BD – Basal Diameter (including the number of samples for mean and maximum measurements).

| Species | Variable |  | n |
| --- | --- | --- | --- |
| Atlantic bluefin tuna | NISP |  | 130 |
|  | Catch |  | 1225 |
|  | TL |  | 68 |
| Gilthead sea bream | NISP |  | 193 |
|  | TL |  | 154 |
| Banded-dye murex | MNI |  | 24 |
| Turbinate monodont | NISP |  | 44 |
|  | MNI |  | 41 |
|  | BD | Mean | 14 |
|  |  | Maximum | 57 |

## Notes

### Competing Interest Statement

The authors have declared no competing interest.

### Summary of Updates

This version of the manuscript has been revised to update the title and include further clarifications on data sources, methodological improvements and adjustments to the scope and focus of the analysis.

